# Integrative genomic strategies applied to a lymphoblast cell line model reveal specific transcriptomic signatures associated with clozapine response

**DOI:** 10.1101/2020.09.22.308262

**Authors:** SAJ de With, APS Ori, T Wang, SL Pulit, E Strengman, J Viana, J Mill, S de Jong, RA Ophoff

## Abstract

Clozapine is an important antipsychotic drug. However, its use is often accompanied by metabolic adverse effects and, in rare instances, agranulocytosis. The molecular mechanisms underlying these adverse events are unclear. To gain more insights into the response to clozapine at the molecular level, we exposed lymphoblastoid cell lines (LCLs) to increasing concentrations of clozapine and measured genome-wide gene expression and DNA methylation profiles. We observed robust and significant changes in gene expression levels due to clozapine (n = 463 genes at FDR < 0.05) affecting cholesterol and cell cycle pathways. At the level of DNA methylation, we find significant changes upstream of the LDL receptor, in addition to global enrichments of regulatory, immune and developmental pathways. By integrating these data with human tissue gene expression levels obtained from the Genotype-Tissue Expression project (GTEx), we identified specific tissues, including liver and several tissues involved in immune, endocrine and metabolic functions, that clozapine treatment may disproportionately affect. Notably, differentially expressed genes were not enriched for genome-wide disease risk of schizophrenia or for known psychotropic drug targets. However, we did observe a nominally significant association of genetic signals related to total cholesterol and low-density lipoprotein levels. Together, these results shed light on the biological mechanisms through which clozapine functions. The observed associations with cholesterol pathways, its genetic architecture and specific tissue effects may be indicative of the metabolic adverse effects observed in clozapine users. LCLs may thus serve as a useful tool to study these molecular mechanisms further.

## Introduction

Antipsychotic drugs (APs) play an important role in the treatment of psychotic disorders such as schizophrenia (SCZ). Clozapine is one of the most effective antipsychotic drugs (AP)(Kane et al., 1988; Leucht et al., 2013; Taylor, 2017). However, the decision to prescribe clozapine is complicated by its potential to induce severe adverse effects(Leucht et al., 2013). The most severe adverse effect, with a prevalence of <1%, is clozapine-induced agranulocytosis, a dramatic reduction of white blood cells(Andersohn et al., 2007). More common adverse effects include weight gain, dyslipidemia and type 2 diabetes. These adverse metabolic effects are, in addition to the chance of developing agranulocytosis, the primary reasons for patient noncompliance and discontinuation of treatment(Cohen, 2014; Weiden et al., 2004).

The biological mechanisms underpinning the effect of clozapine, as well as its adverse effects, remain elusive. A twin study estimated that the heritability of APs-induced weight gain is approximately 60%(Gebhardt et al., 2010), suggesting a substantial role for genetic factors. Candidate genes studies of clozapine-induced adverse effects have yielded ambiguous results and lack consistent replication (reviewed by(Chowdhury et al., 2011; Lett et al., 2012; Müller et al., 2013; Roerig et al., 2011; Yan et al., 2013)). Two genome-wide association studies (GWAS) investigating antipsychotic-induced metabolic adverse effects have yielded inconclusive findings, primarily due to insufficient sample sizes and the potential polygenic nature of this trait(Adkins et al., 2011; Malhotra et al., 2012).

Intermediate molecular phenotypes, such as gene expression studies in specific cell lines or tissues, may improve our understanding of the molecular function of clozapine. Previous studies have found that atypical antipsychotic drugs may induce cholesterol metabolism through transcription factors such as sterol regulatory element binding proteins (*SREBP1* and *2*)(Ferno et al., 2011), suggesting that drug-induced cholesterol metabolism is related to these adverse metabolic effects. However, such findings were not consistently replicated when profiling whole blood(Ferno et al., 2011; Harrison et al., 2016; Vik-Mo et al., 2008). It is possible that changes in DNA methylation could mediate changes in gene expression, such as the AP-induced hypomethylation of the *FAR2* gene leading to insulin resistance(Burghardt et al., 2016). However, such studies are currently limited and those performed provide inconsistent results(Burghardt et al., 2016; Houtepen et al., 2016; Kinoshita et al., 2017; Melas et al., 2012; Ota et al., 2014; Rukova et al., 2014; Stapel et al., 2017; Swathy et al., 2017; Swathy and Banerjee, 2017).

A major obstacle towards understanding clozapine-induced metabolic effects is that clozapine therapy is a relatively rare (∼6%) treatment plan in patients diagnosed with schizophrenia(Burghardt et al., 2016; Stroup et al., 2016), of which then only 1% develop CIA. Such factors challenge our ability to adequately sample a large and controlled prospective cohort of patients thereby limiting progress in understanding both metabolic and hematological adverse effects. To augment the lack of available *in vivo* data, we implemented an *in vitro* lymphoblast cell line (LCL) model to study the effects of drug exposure at the molecular level. Cell-based models have been successfully employed for pharmacogenomic studies, including LCL models to study clozapine function(de With et al., 2015; Morag et al., 2010; Welsh et al., 2009; Wen et al., 2012). Here, we exposed LCLs to increasing doses of clozapine and collected both expression and methylation profiles. Through integrative genomic analyses, we aim to extrapolate *in vitro* molecular signatures of clozapine towards relevance of *in vivo* function and study clozapine response without the need to assemble a large cohort of patients. We identified significant changes in transcriptomic signatures associated with cholesterol and cell cycle pathways. By then integrating these molecular profiles with genome-wide association study (GWAS) summary statistics of different traits and diseases, we show that clozapine-associated genes also overlap with genetic signals related to total cholesterol and low-density lipoprotein (LDL) levels but not schizophrenia genetic risk. Clozapine-associated genes are furthermore related to specific human tissues, such as liver and those involved in immune, endocrine and metabolic functioning.

## Material and Methods

### Lymphoblast cell lines

We used lymphoblast cell lines (LCLs) from four unrelated samples, all part of the collection of Utah residents of Northern and Western European ancestry (HapMap CEPH/CEU phase 1)(Consortium and †The International HapMap Consortium, 2003). We obtained LCLs from the Coriell Institute for Medical Research (Camden, NJ, USA) and maintained the cell lines as previously described(Consortium and †The International HapMap Consortium, 2003; de With et al., 2015). To study methylation changes after exposure to clozapine, we performed a separate experiment using six LCLs (HapMap, CEPH/CEU phase 1) consisting of two parent-offspring trios.

### In vitro experimental design and clozapine exposure

We exposed LCLs to different clozapine concentrations, with clinical concentration set at 2µM(Baumann et al., 2004). Clozapine, purchased from Sigma Aldrich, was dissolved in culture medium with dimethyl sulfoxide (DMSO), with a maximum concentration of 0.1%. Cell lines were exposed for 24 hours to clinical concentration (Supplementary Methods), 10x, 50x and 100x clinical concentration (20µM-100 µM-200 µM clozapine) and vehicle (DMSO); each concentration was measured in 4 cell lines, after which RNA was obtained for gene expression analysis (Supplemental Figure 1A). To study DNA methylation changes in response to clozapine, we performed an experiment similar to the gene expression study; LCLs were exposed to different concentrations of clozapine (vehicle (DMSO), 1x, 20x, 40x and 60 times clinical concentration) and exposure times were 24h and 96h (Supplemental Figure 1B). We measured cell viability using the TC10 automated cell counter.

**Figure 1.**
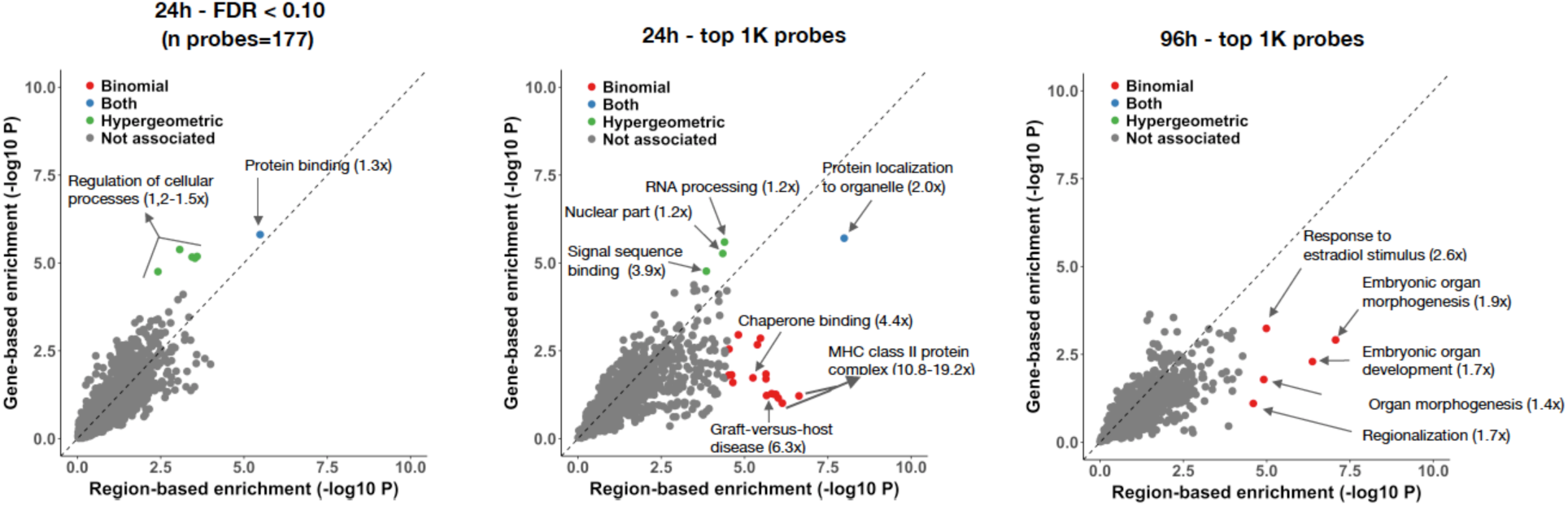
Clozapine-associated DNAm probes concentrate to specific biological annotations. Genomic Regions Enrichment of Annotations Tool (GREAT) was used to assign biological context to genomic regions associated to clozapine exposure. GREAT uses the annotation of nearby genes and regulatory elements to performed enrichment analysis across known functional databases. The x-axis shows the -log 10 p-value of the region-based analysis (binomial test), while the y-axis shows the enrichment for the gene-based analysis (hypergeometric overlap test). Results are shown for (A) the top probes after 24 hours (FDR < 0.10, n=177), (B) the top 1,000 probes after 24 hours, and (C) the top 1,000 probes after 96 hours. Each dot represents an annotation. Significant annotations, after Bonferroni correction, are color-coded according to the test used.

### Sample processing and data collection

After desired exposure time, we lysed cells and performed RNA and DNA collections using column-based extraction methods from Qiagen according to manufacturer’s instructions (Supplementary Methods). Gene expression profiling was carried out using Illumina® HumanHT-12 v4 Expression BeadChip technology. DNA methylation assays were performed with Illumina® Infinium HumanMethylation450 Beadchip arrays.

### Data preprocessing and normalization

We processed raw gene expression values using the “Lumi” R-package(Du et al., 2008). We log2 transformed and quantile normalized the raw data, keeping only expressed gene transcripts (detection p < 0.01) for further analysis (22,926 probes). We processed DNA methylation values using the “WateRmelon” Bioconductor package(Pidsley et al., 2013), removing probes that were known to cross-hybridize, probes containing SNPs in target CpG regions, probes with detection p-value greater than 0.01 in 5% of samples, and probes with beadcounts > 3 (n=86,068 probes in total)(Chen et al., 2013; Price et al., 2013). We normalized the data using the *dasen* function and computed β-values, defined as the ratio of the methylated probe intensity and the overall intensity (sum of methylated and unmethylated probe intensities), to measure methylation levels. To limit the effect of heteroskedasticity, we included only variable probes with β-values between 0.2 – 0.8 in our analyses (165,014 probes)(Du et al., 2010).

### Statistical analyses

To detect clozapine-induced molecular changes, for each probe, we tested for association between gene expression (1) or DNA methylation levels (2) with increasing clozapine concentrations using the following linear models implemented in R using the “Limma” package(Smyth, n.d.):

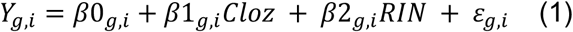

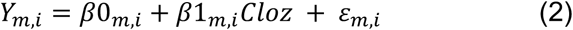

where Y is the normalized gene expression (g) or DNA methylation levels (m) for an individual probe, b0 the intercept, b1 the effect of clozapine concentration, b2 the effect of RIN, and e the residual variation of the model. We ran each model for each probe per individual i and subsequently performed a meta-analysis across all individuals by combining p-values using Stouffer’s method with directionality of the effect sizes taken into account. We included RNA-integrity number (RIN) as a covariate in the gene expression model and applied a Bonferroni correction to correct for multiple testing, resulting in a significance threshold of p < 2.18×10^−6^ for the gene expression analysis (n = 22,926 probes) and p < 3.03×10^−7^ for the DNA methylation analysis (n = 165,014 probes).

### Gene ontology analysis

We performed functional gene ontology analysis using DAVID (Database for Annotation, Visualization and Integrated Discovery, version 6.8, interrogated February 2018)(Huang et al., 2009a, 2009b), with default settings (Supplementary Methods).

### Functional enrichment analysis of DNA methylation data

We used Genomic Regions Enrichment of Annotations Tools (GREAT, v3.0) to predict the biological function of the top methylation probes associated with clozapine exposure. GREAT links both proximal and distal genomic CpG sites with their putative target genes and implements both a gene-based test and a region-based test using the hypergeometric and binomial test, respectively, which allows us to assess enrichment of genomic regions in biological annotations of pathway databases (Supplementary Methods)(McLean et al., 2010). The statistical outputs of GREAT for both gene-based and region-based tests were subsequently adjusted for multiple testing using Bonferroni correction.

### Weighted gene co-expression network analysis

We performed a gene expression network analysis using weighted gene co-expression network analysis (WGCNA) in R. Briefly, WCGNA identifies distinct modules using the shared variation in gene expression based on pairwise correlation. To account for the biases related to differing probe numbers between genes assayed on the array, we provided as input the mean probe expression of genes residing within nominally significant differentially expressed genes (p<0.05), considering 5,708 probes within 4,897 genes(Horvath, 2011; Langfelder and Horvath, 2008; Zhang and Horvath, 2005). To assign biological function to each WCGNA module, we performed gene ontology analysis using DAVID we performed.

### Additional DNA methylation analyses

We ran a candidate gene study for CpG sites in close proximity of genes with evidence of clozapine-induced differential gene expression. Methylation probes within the gene body, in the 3’ and 5’ untranslated regions and up to 1,500 nucleotides upstream of the transcription start site of the 463 ‘top genes’ (p<0.05) were selected for a post-hoc analysis (n = 1,004). These results can be found in Table 1 of the Supplemental Material.

**Table 1.**
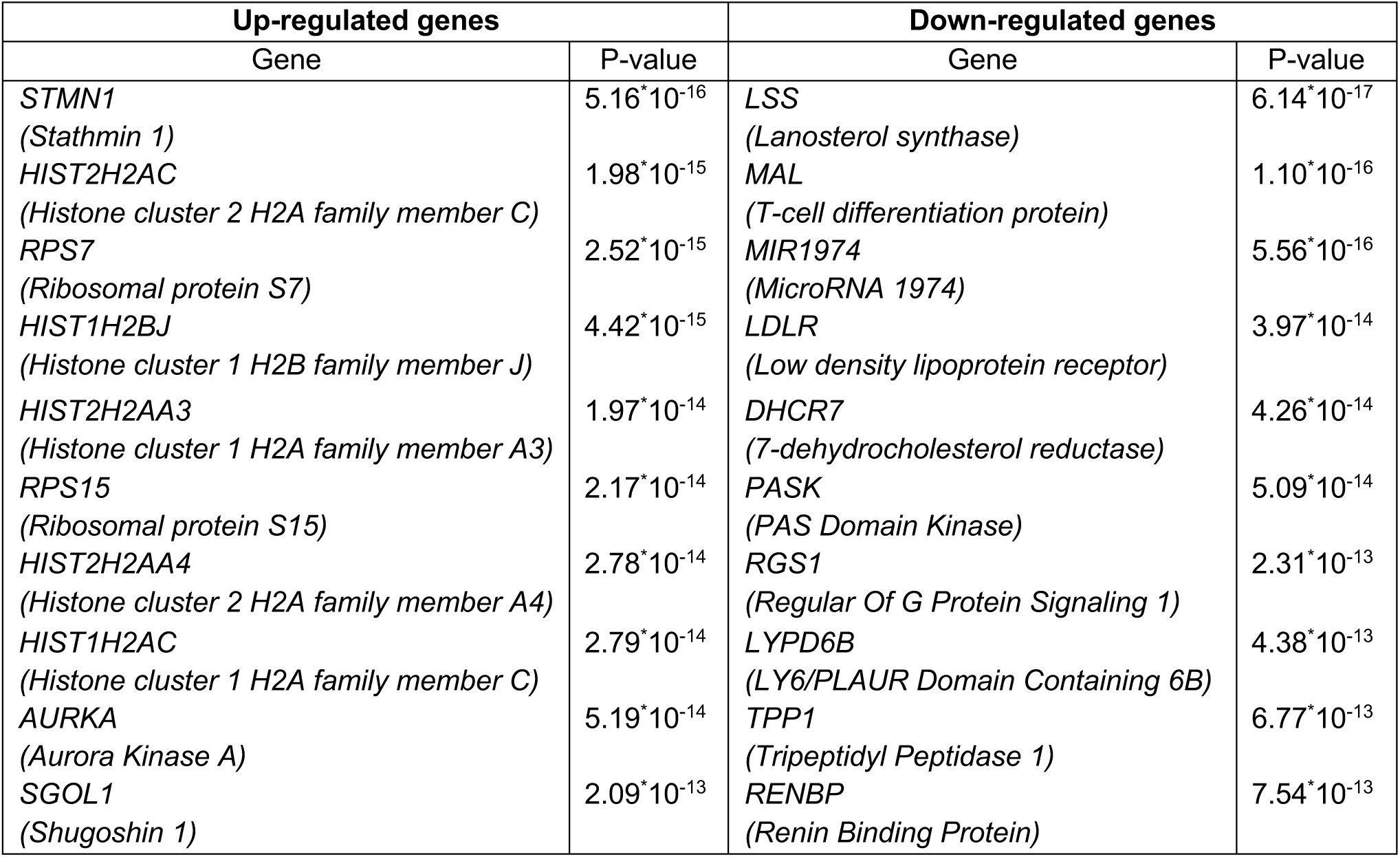
Top 10 up- and downregulated genes after clozapine exposure. Gene symbols are shown with corresponding gene name and p-value of differential gene expression analysis. A full list of genes is in Supplementary Table 1.

| Up-regulated genes |  | Down-regulated genes |  |
| --- | --- | --- | --- |
| Gene | P-value | Gene | P-value |
| *STMN1*<br>(Stathmin 1) | $5.16 \times 10^{-16}$ | *LSS*<br>(Lanosterol synthase) | $6.14 \times 10^{-17}$ |
| *HIST2H2AC*<br>(Histone cluster 2 H2A family member C) | $1.98 \times 10^{-15}$ | *MAL*<br>(T-cell differentiation protein) | $1.10 \times 10^{-16}$ |
| *RPS7*<br>(Ribosomal protein S7) | $2.52 \times 10^{-15}$ | *MIR1974*<br>(MicroRNA 1974) | $5.56 \times 10^{-16}$ |
| *HIST1H2BJ*<br>(Histone cluster 1 H2B family member J) | $4.42 \times 10^{-15}$ | *LDLR*<br>(Low density lipoprotein receptor) | $3.97 \times 10^{-14}$ |
| *HIST2H2AA3*<br>(Histone cluster 1 H2A family member A3) | $1.97 \times 10^{-14}$ | *DHCR7*<br>(7-dehydrocholesterol reductase) | $4.26 \times 10^{-14}$ |
| *RPS15*<br>(Ribosomal protein S15) | $2.17 \times 10^{-14}$ | *PASK*<br>(PAS Domain Kinase) | $5.09 \times 10^{-14}$ |
| *HIST2H2AA4*<br>(Histone cluster 2 H2A family member A4) | $2.78 \times 10^{-14}$ | *RGS1*<br>(Regular Of G Protein Signaling 1) | $2.31 \times 10^{-13}$ |
| *HIST1H2AC*<br>(Histone cluster 1 H2A family member C) | $2.79 \times 10^{-14}$ | *LYPD6B*<br>(LY6/PLAUR Domain Containing 6B) | $4.38 \times 10^{-13}$ |
| *AURKA*<br>(Aurora Kinase A) | $5.19 \times 10^{-14}$ | *TPP1*<br>(Tripeptidyl Peptidase 1) | $6.77 \times 10^{-13}$ |
| *SGOL1*<br>(Shugoshin 1) | $2.09 \times 10^{-13}$ | *RENBP*<br>(Renin Binding Protein) | $7.54 \times 10^{-13}$ |

### GTEx cross tissue analysis

To translate the *in vitro* effects of clozapine to *in vivo* human biology, we investigated how clozapine genes behave across 22 human tissues represented in the GTEx data set. We downloaded gene level quantifications (version 6, date: April 24, 2019) from the GTEx Project web portal(Melé et al., 2015) and transformed gene expression values using a log2 transformation (1 + RPKM value). We then tested 1) if clozapine-associated genes have higher or lower average expression within each tissue and 2) if between tissue distance is different for clozapine-associated genes compared to the expected based on chance. To calculate between tissue distance, we used the top half most variable genes across GTEx samples that were also significantly detected in our *in vitro* assay (n=7,025). We then performed multidimensional scaling using the isoMDS() function in the MASS R package (v7.3)(Melé et al., 2015; Venables and Ripley, 2002) and calculated between tissue Euclidean distances using the dist() function in R (v.3.3.3) and the median gene expression values for each tissue for clozapine associated genes (N=463). To established whether clozapine-associated genes significantly deviated from the expected, we established a null distribution by repeating this procedure using the Euclidean distance for samplings of similarly-sized sets of genes. Within each sampling, genes were sampled with a probability that matched the distribution of average gene expression of clozapine-associated genes in our LCL experiment. This was done to account for differentially expressed genes having higher gene expression levels. Our null distribution therefore better captured the expected effect. Null distributions were built separately for the expected average gene expression within each tissue and for the expected Euclid distance for each tissue pair. P-values were then established by calculating the proportion of samplings that were higher or lower than the statistic of the observed clozapine-associated genes and corrected for multiple testing by Bonferroni correction. Analyses were conducted separately for genes upregulated and downregulated after clozapine exposure.

### MAGMA gene-set analysis

To investigate if clozapine-associated genes were enriched for schizophrenia or cardiovascular-related genetic association signals, we performed gene-set analysis using MAGMA (multimarker analysis of genomic annotation)(de Leeuw et al., 2015). We tested the following three gene-sets, each with three different significance thresholds:

1. All genes differentially expressed, either up- or downregulated;
2. Upregulated differentially expressed genes;
3. Downregulated differentially expressed genes.

We obtained GWAS summary statistics for schizophrenia(Schizophrenia Working Group of the Psychiatric Genomics Consortium, 2014), body mass index(Yengo et al., n.d.), coronary artery disease(Nikpay et al., 2015), type 2 diabetes(Scott et al., 2017), total cholesterol, triglycerides, high-density-lipoprotein, and low-density-lipoprotein(Scott et al., 2017; Surakka et al., 2015). For each GWAS, we computed aggregate gene-level test statistics using a 10kb window around the transcription start and end site of each gene; a total of 11,533 genes were available for analysis. We then tested for association between GWAS gene level Z-scores and experimental gene-sets (defined above). We estimated linkage disequilibrium (LD) using the 1000 Genomes European reference panel(1000 Genomes Project Consortium et al., 2015; Scott et al., 2017; Surakka et al., 2015) and ran MAGMA using a two-sided competitive test by accounting for probes tested in our experiment while also correcting for gene size, SNP density, minor allele count and LD.

### Antipsychotic drug gene-sets and enrichment of schizophrenia heritability

As a complementary analysis, we investigated whether manually curated antipsychotic drug gene-sets overall are associated with schizophrenia heritability. We performed gene-set analysis on antipsychotic drug targets that were previously reported to be enriched for schizophrenia heritability [55]. Drug target list were curated using drug-gene interaction databases(Roth et al., 2000; Wagner et al., 2016). We identified 50 sets of drugs with ATC (Anatomical Therapeutic Chemical) code N05A, a drug class to which all antipsychotic drugs belong, including clozapine. Using the schizophrenia GWAS as input to MAGMA, we ran gene-set analysis (1) per individual N05A antipsychotic drug gene-set, (2) for all N05A drug target genes combined, and for (3) all N05A drug target genes combined excluding clozapine.

### LD Score Regression estimation of heritability and genetic correlations

Pre-computed LD scores from the 1000 Genomes European reference panel provided through the LD Score Regression (LDSR) *github* and GWAS summary statistics file processed and reformatted to s*umstats* format were used as input to (LDSR) to estimate SNP-based heritability (SNP-h^2^)(B. K. Bulik-Sullivan et al., 2015). Genetic correlations between traits were estimated using cross-trait LDSR via the *--rg flag(B. Bulik-Sullivan et al*., *2015)*.

## Results

### Clozapine exposure induces widespread gene expression changes

We exposed lymphoblast cell lines to different concentrations of clozapine and tested for subsequent changes in gene expression levels (Supplemental Figure 1A). In total, we tested 22,926 gene expression probes, of which 5,708 showed nominal significance (p<0.05) and 518 probes exceeded a Bonferroni-corrected p < 2.18^*^10^−6^. These 518 probes consisted of 234 up-regulated probes and 284 down-regulated probes, representing a total of 463 unique genes. The top 10 up-regulated and down-regulated probes are shown in Table 1. See Supplement for the complete list of differentially expressed genes.

Gene ontology enrichment analysis of up-regulated transcripts showed significant functional enrichment for cholesterol metabolism (13 genes, p = 4.15^*^10^−15^) and steroid biosynthesis (11 genes, p = 1.01^*^10^−8^) (Table 2), while analysis of down-regulated transcripts were enriched for cell division processes and related annotations, such as mitosis (44 genes, p = 1.87^*^10^−39^), chromosome (49 genes p = 3.03^*^10^−35^) and nucleosome (16 genes, p = 3.12^*^10^−14^) and other cell cycle pathways (Table 2).

**Table 2.** Main functional annotations associated with clozapine exposure based on gene expression. Output of pathway enrichment analyses using DAVID is shown. The enrichment score is used as the main metric of importance. It is defined as the geometric mean of all enrichment p-values of each annotation term within the group. It is expressed as the minus log of the p-value, an enrichment score of 1.3 is nominally significant. This table shows clusters with an enrichment score > 4.6, corresponding to a p-value < 0.01. Fold enrichment is a measure to express the enrichment of this particular group of genes in comparison with the genes in the human genome. A list of all pathways is available in supplemental information.

| Functional annotation cluster | Enrichment score | Best P-value | Best fold enrichment | Average P-value | Average fold enrichment | Number of pathways |
| --- | --- | --- | --- | --- | --- | --- |
| *Upregulated pathways* |  |  |  |  |  |  |
| Cholesterol/lipid/steroid biosynthesis | 8.44 | $4.15 \times 10^{-15}$ | 46.92 | $1.19 \times 10^{-3}$ | 18.46 | 19 |
| Cholesterol/Steroid biosynthesis | 4.77 | $1.01 \times 10^{-8}$ | 68.45 | $4.66 \times 10^{-4}$ | 54.32 | 3 |
| *Down-regulated pathways* |  |  |  |  |  |  |
| Mitosis/cell division | 31.26 | $1.87 \times 10^{-37}$ | 14.77 | $1.12 \times 10^{-21}$ | 10.69 | 5 |
| Chromosome/centromere | 16.47 | $3.03 \times 10^{-35}$ | 21.32 | $7.76 \times 10^{-8}$ | 15.79 | 9 |
| Histone/nucleosome | 6.93 | $3.12 \times 10^{-14}$ | 20.72 | $1.73 \times 10^{-3}$ | 9.13 | 18 |
| Spindle | 6.02 | $4.95 \times 10^{-12}$ | 12.89 | $7.45 \times 10^{-4}$ | 10.13 | 6 |
| Nucleotide/ATP-binding | 5.72 | $1.46 \times 10^{-13}$ | 2.71 | $7.78 \times 10^{-5}$ | 2.47 | 5 |
| Microtubule/kinesin | 4.60 | $1.90 \times 10^{-4}$ | 16.92 | $4.99 \times 10^{-2}$ | 8.56 | 19 |

### Gene expression network analysis after clozapine exposure

WGCNA network analysis of clozapine-induced differentially expressed genes (N=4,987) yielded 15 co-expression modules, ranging in size from n=61 to 1,791 genes. Five gene co-expression modules were altered upon clozapine exposure: M14 (upregulated, 61 genes), M10 (upregulated, 155 genes), M9 (upregulated, 158 genes), M3 (upregulated, 579) and M1 (downregulated, 1,791 genes), which were nominally significant. The M14 co-expression module was enriched for genes involved in cholesterol metabolism (13 genes, p = 4.8^*^10^−15^), the M1 co-expression module was enriched for genes involved in cell cycle (133 genes, p = 7.3^*^10^−32^) and the M9 and M3 co-expression module was enriched for mitochondrial genes (15 genes, p = 6.7^*^10^−6^ and 46 genes p = 3.8^*^10^−7^ respectively). The M10 co-expression module was enriched for genes involved in the nucleosome (9 genes, p = 1.4^*^10^−7^).

### Minimal changes in DNA methylation at single CpG sites after clozapine exposure

We then performed DNA methylation profiling to assess whether these effects were due to epigenetic changes. For statistical analysis, we applied an analytical approach similar to our gene expression analyses. Targeted analysis on the 1,004 CpG sites near the 463 differentially-expressed genes revealed 3 probes exhibiting significant changes in DNA methylation (p < 1.08^*^10^−4^), including a probe upstream of the low-density lipoprotein receptor (LDL-R) gene (cg22971501, p=4.75*10^−5^) after 24h; one probe upstream of the cyclin F (CCNF) gene showed a significant change in DNA methylation after 96h (Table S3). Beyond these examples, global methylation differences were not observed after 24h or 96h at the level of individual probes.

### DNA methylation is affected at the pathway level

To investigate if our top associated DNAm probes aggregated to changes at the pathway level, we performed pathway enrichment analysis using GREAT, which incorporates functional annotation from various databases to predict cis-regulatory function of genomic regions of interest. When considering probes exhibiting FDR < 10%, we observed enrichment for protein binding and regulation of cellular processes after 24h of clozapine treatment. When considering the top 1000 probes, we also observed enrichment of immune-related functions, such as the Major Histocompatibility Complex (MHC) class II protein complex and Graft-versus-host disease after 24h. We found significant enrichment for estradiol regulation and various embryonic developmental processes (Figure 1 and Table S5) when considering the top 1000 probes after 96h.

### DNA methylation changes are time-dependent

To examine if clozapine affected each time point similarly, we correlated effect sizes at each CpG probe between time points. Across all probes, we found a significant negative correlation between the Z-score of the association between DNA methylation levels and clozapine exposure at 24h and 96h (Pearson r = -0.32, P<2.2×10^−16^). This correlation was preserved among probes at FDR < 10% at 24h (n=177, Pearson r = -0.38, P=1.3×10^−7^) but not among the top probes at 96 hours (n=177, Pearson r = -0.12, P = 0.10), highlighting possible time-dependent effects after clozapine exposure (Figure 2).

**Figure 2.**
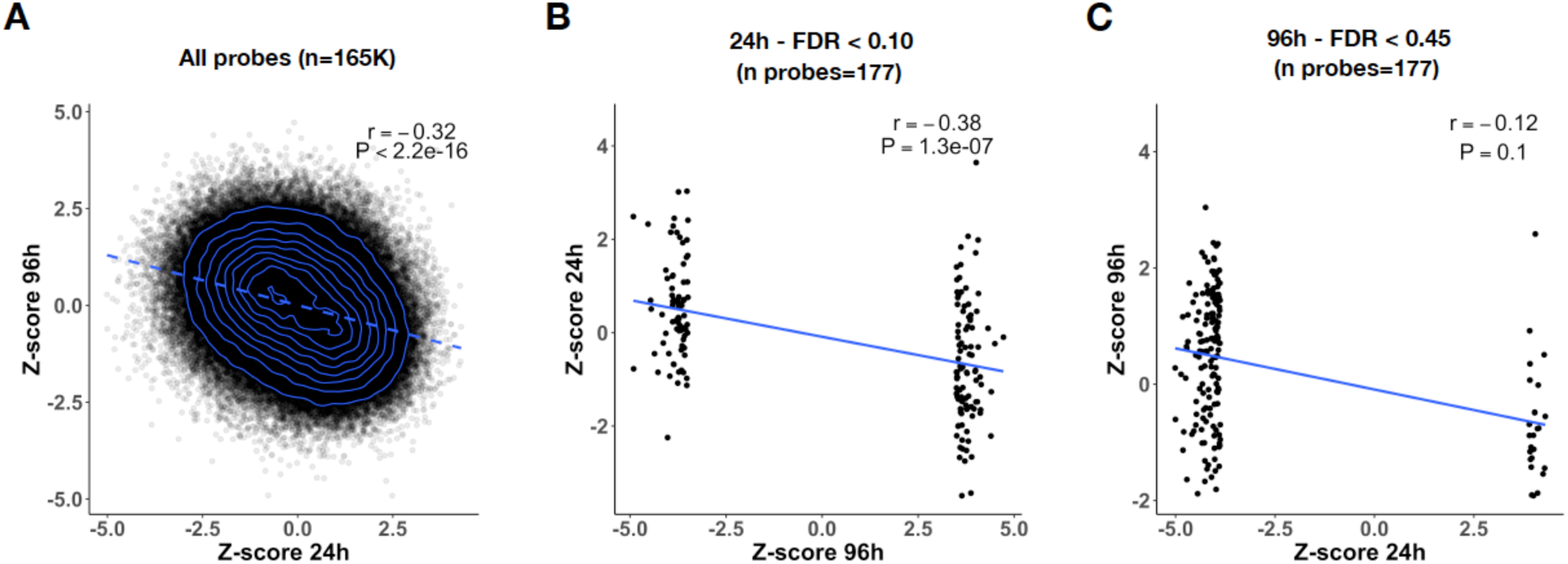
Clozapine-induced DNA methylation changes are time-dependent. Correlation analysis was performed between DNAm probe associations after 24 and 96 hours of clozapine exposure. The Pearson correlation was computed for (A) all probes tested, (B) the top probes associated after 24hour exposure (FDR < 0.10, n = 177 probes), and (C) the top 177 probes after 96 hours of drug exposure.

### Clozapine transcriptomic profiles highlight multi-tissue effects in GTEx

We then asked whether our *in vitro* derived transcriptomic signatures could be used to help translate the function of clozapine in humans. For this purpose, we used gene expression data from the GTEx Project, including LCLs (n = 22 GTEx tissues, Figure 3A). First, we overlapped preferentially-expressed genes of each GTEx tissue, as previously reported(Melé et al., 2015), with the clozapine-associated genes detected in our assay. Preferentially expressed genes in GTEx-LCLs exhibited the most overlap with the genes identified by our experiment followed by whole blood (Supplementary Figure 2). We then investigated if the mean tissue expression of clozapine-associated genes was significantly different from the mean expression of all the genes tested in our experiment. We found that genes downregulated by clozapine have lower expression in all tissues except testis and LCLs (Figure 3B and Table S6). Downregulated genes are enriched for cell cycle processes and their higher expression in the testis and LCLs likely represent the proliferative nature of these tissues. Genes upregulated by clozapine have significantly higher average expression in liver, muscle, lung, and fibroblasts, and lower average expression in testis tissue. We then asked whether clozapine-associated genes have different between-tissue distances as a proxy to investigate possible functional links with other tissues. Distance is calculated using the Euclidean distance measure. A lower value indicates that two tissues are more similar while a higher distance value indicates that these tissues are more dissimilar. Of the 406 tissue pairs, we found that genes upregulated by clozapine have significantly deviating between-tissue distances across 31 pairs of tissues (Figure 3C and Table S7). Spleen tissue stands out with having the most different distance with other tissues (N=20). We also found multiple differences for adipose, lung, and breast tissue pairs. The distance between cervix uteri and ovary tissue, adrenal gland and thyroid tissue, muscle and nerve tissue are significantly more dissimilar as well for upregulated clozapine genes compared to all genes detected in our assay. For downregulated clozapine genes, we detected significantly dissimilar distances for 27 tissue-pairs, all involving testis tissue (Figure 3C).

**Figure 3.**
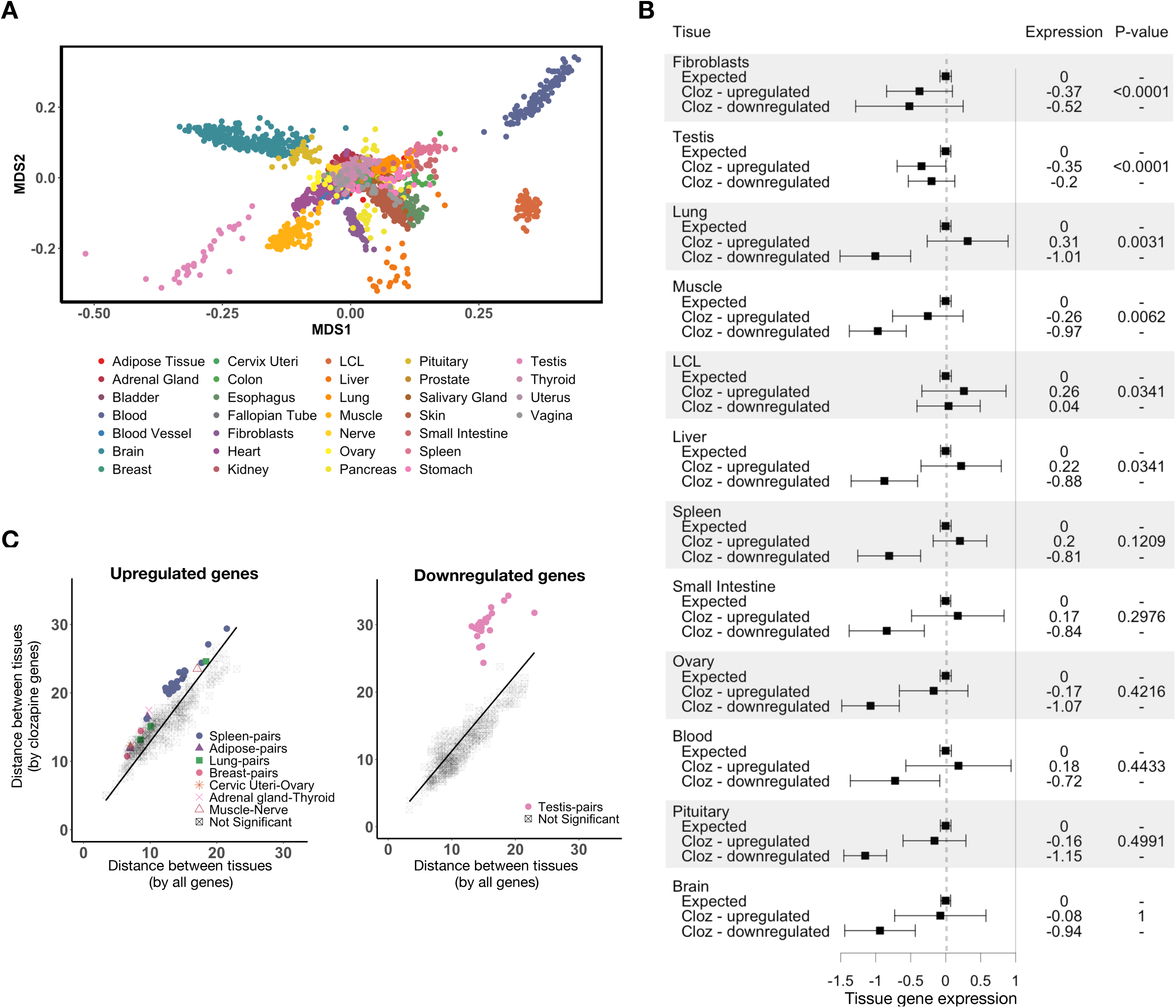
Clozapine-associated genes show tissue-specific expression patterns in GTEx. (A) Using gene expression data from the GTEx project, multidimensional scaling (MDS) was performed to visualize tissues and their relationships in a parsimonious way. The top half most variable genes (n=7,025), that were also significantly detected in our experiment, across a random subset of 2,000 GTEx samples were used as input in the MDS analysis. (B) A forest plot visualizing mean tissue gene expression of clozapine-associated genes across several tissues. Clozapine genes were divided into groups that are up regulated and down regulated after drug exposure. The expected mean gene expression, based on 10,000 weighted samplings, is shown as well (dotted vertical line). Within each tissue, the observed mean tissue gene expression (x-axis) is normalized by subtracting the expected mean gene expression. P-values indicate whether the mean gene expression of clozapine up regulated genes significantly deviate from the expected mean expression within a tissue. P-values are corrected for the number of tissues tested. Downregulated genes were not tested. See Supplementary Tables for results of all tissues (C) Between tissue distance across all GTEx tissue pairs. Each point represents one tissue pair. The y-axis shows the Euclidean distance between tissues computed using only clozapine-associated genes. The x-axis shows the expected mean Euclidean distance across 50,000 weighted samplings. Tissue pairs for which the between tissue distance, based on clozapine genes only, significantly deviates from mean expected distance (Padjusted <0.05), are color-coded. P-values are adjusted for multiple testing by Bonferroni correction (n test = 465 pairs x tests = 930).

### Clozapine transcriptome signatures are not enriched for schizophrenia disease risk

Previous studies have found associations between schizophrenia genetic susceptibility and antipsychotic drug targets(Gaspar and Breen, 2017; Skene et al., 2018). To investigate whether clozapine-induced *in vitro* gene expression signatures are also associated with genetic susceptibility of schizophrenia, we conducted gene-set enrichment analysis using all differentially expressed genes (n=463). We did not observe a significant enrichment (p = 0.91). We then considered upregulated genes, and did not observe significant enrichment (p = 0.74), nor for downregulated genes (p = 0.64). Gene sets defined according to <1% and <5% FDR did not change these findings (Table S8).

To further understand these findings, we investigated whether SCZ genetic susceptibility aggregates to antipsychotic drug target genes with and without clozapine targets. As SCZ genetic risk has been associated with antipsychotic drug target genes, it could be that this signal is primarily driven by non-clozapine genes. To examine this, we used drug target gene lists from drug-gene interaction databases as previously reported(Roth et al., 2000; Wagner et al., 2016). We were able to extract 53 drug target gene-sets belonging to the N05A class of drugs with antipsychotic actions, including clozapine. Across drug target gene-sets, we mapped all targets to 104 unique genes of which 41 were detectable in our gene expression data but none overlapped with differentially expressed genes identified. We found no evidence for SCZ risk to be enriched in antipsychotic drug target genes overall (n = 96 genes, p = 0.96) nor with clozapine targets excluded (n = 52, genes, p = 0.60). We did observe a strong concordance between the p-values of individual drug gene-sets tested in our analysis and the p-values reported by the previous study(Gaspar and Breen, 2017) (n = 50 drug gene sets, rho = 0.85, p = 1.49^*^10^−15^), indicating our analysis framework was able to reproduce previous findings at the level of individual drug target gene-sets. Our analysis however does not observe an association between schizophrenia genetic risk and clozapine-associated genes nor antipsychotic drug target genes.

### Genes upregulated after clozapine exposure show an association with GWAS signal of total cholesterol and low-density lipoprotein

Next, we explored whether differentially expressed genes were enriched for genetic signal of cholesterol-and cardiovascular disease-related traits. Using summary statistics of large GWASs for each trait, we observe significant SNP-h^2^ and a rich correlation structure between these traits (Figure 4A and 4B). We then integrated the observed SNP-h2 with our detected clozapine transcriptomic signatures. While no association remained significant after correction of multiple testing (72 tests, p < 6.9^*^10^−4^), we did observe an increasing association between total cholesterol and LDL heritability and clozapine gene-sets across more stringent thresholds of differentially expressed genes (i.e. <FDR 5%, <FDR 1%, and Bonferroni correction (Figure 4C and Table S9). Such trends were not observed for HDL or any of the other traits tested, including SCZ.

**Figure 4.**
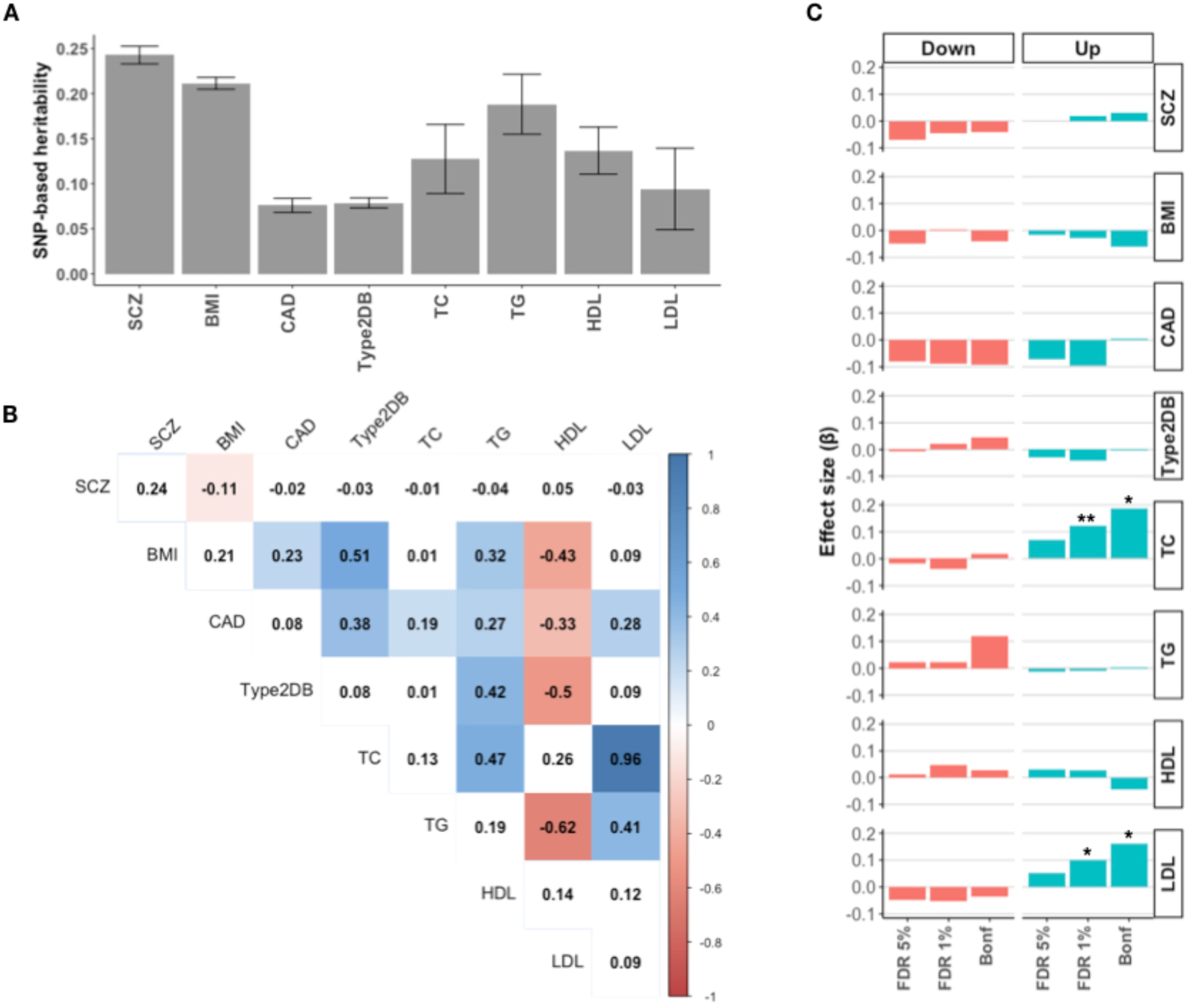
Clozapine gene-set analysis across cardiovascular traits. (A) SNP-based heritability estimates on the observed scale. (B) Genetic correlations between traits used in the analysis. The magnitude of the genetic correlation is only color-coded for estimates with p-value < 0.01. (C) MAGMA effect sizes (β) of GWAS trait associations with clozapine target genes are shown for down-and unregulated genes identified to be differentially expressed at FDR 5%, FDR 1%, and Bonferroni correction thresholds. Asterisks denote nominal significance of *P<0.05, **P<0.01. SCZ = schizophrenia, BMI = body mass index, CAD = coronary artery disease, Type2DB = type 2 diabetes, TC = total cholesterol, TG = triglycerides, HDL = high-density-lipoprotein, LDL = low-density-lipoprotein.

## Discussion

We used an *in vitro* cell system of LCLs to study molecular effects of clozapine exposure to better understand the mechanisms involved in adverse effects of psychotropic drug use and make several important observations. First, genome-wide gene expression profiling of cells exposed to clozapine showed strong activation of cholesterol metabolism and deactivation of genes involved in cell cycle processes. These findings align with our observation of changed levels of DNAm upstream of the LDL-R and CCNF genes after clozapine exposure. Second, DNAm analyses suggest that effects of clozapine are time-dependent indicating that time of drug exposure is an important experimental variable to take into account when studying clozapine. Third, integration of *in vitro* transcriptomic signatures with human tissues in GTEx highlight liver tissue and several immune and endocrine tissues as possible downstream effectors of clozapine exposure. Finally, genetic analysis of the results suggests that clozapine-response in LCLs is independent from schizophrenia disease risk and depleted from known drug targets of antipsychotic drugs, while it is likely linked to the genetic architecture involved in cholesterol and low-density lipoprotein levels.

*In vitro* gene expression changes after exposure to clozapine have been reported before for various cell lines. Here, we for the first time applied a genome-wide analysis of gene expression in lymphoblastoid cell lines in response to clozapine exposure. The central role of cholesterol metabolism in the response to clozapine is concordant with previously reported studies performed in other cell types and for different antipsychotic drugs(Ferno et al., 2011; Foley and Mackinnon, 2014). Even though most of these were based on candidate genes rather than by genome-wide analyses, the results consistently implicate cholesterol metabolism in response to antipsychotic drugs (Choi et al., 2009; Fernø et al., 2009, 2005; Hu et al., 2010; Lauressergues et al., 2012, 2011, 2010; Liu et al., 2009; Raeder et al., 2006; Vik-Mo et al., 2009; Yang et al., 2007, 2009). At the gene network level, clozapine-associated genes group to clear gene clusters. In addition to cholesterol metabolism, we observed mitochondrial and nucleosome pathways to be upregulated after clozapine exposure, while cell cycle processes were downregulated, for example.

A small number of studies have identified a link between genetic markers in genes involved in cholesterol metabolism and adverse effects of antipsychotic treatment, helping to further substantiate these findings (Chowdhury et al., 2011; Yang et al., 2016, 2015). Additional studies suggest that alterations in cholesterol metabolism by antipsychotic drugs may contribute to the beneficial effects in treating psychosis. Cholesterol is extremely important in brain development and in sustaining neuronal connections and myelination(Dietschy, 2009). Furthermore, Le Hellard et al. described an association between genes important in cholesterol metabolism (*SREBP1* and *SREBP2)* and schizophrenia(Dietschy, 2009; Le Hellard et al., 2010), which was confirmed in a schizophrenia GWAS(Schizophrenia Working Group of the Psychiatric Genomics Consortium, 2014).

In addition to significantly upregulated genes, we also identified 284 genes that are downregulated in the presence of clozapine, with significant enrichment for genes involved in cell cycle pathways. A small number of studies have shown differing gene expression levels in cell cycle genes in patients with schizophrenia, compared to healthy controls (Gassó et al., 2017; Lin et al., 2016; Okazaki et al., 2016; Wang et al., 2010; Yang et al., 2017). A similar connection with antipsychotic drug effects has not been reported before. It is possible that genes involved in cell cycle play a role in the etiology of schizophrenia, making it a possible target of antipsychotic drugs as clozapine. Conversely, the described effect may be a direct consequence of clozapine toxicity. We have shown before that clozapine, in high concentrations, has a direct effect on viability of lymphoblastoid cell lines (de With et al., 2015). Downregulation of genes involved in cell cycle processes could thus be a direct (toxic) effect of clozapine. The toxic effects of clozapine and its metabolites have been associated with clozapine-induced agranulocytosis(de With et al., 2015; Lahdelma et al., 2010; Pereira and Dean, 2006; Williams et al., 1997). We note that for this study, we used LCLs, which are blood-derived cells, but from a different progenitor cell and with different cellular functions than neutrophils. The question remains whether findings from *in vitro* LCL can be directly extrapolated to *in vivo* effects of clozapine on neutrophils.

While we found strong effects of clozapine exposure at the gene expression level, we did not observe similar global effects at the level of DNA methylation. We did find 3 differently methylated CpGs located within genes implicated by our gene expression analysis, including the LDL receptor gene. These effects, however, were not observed after 96h of clozapine exposure. Additionally, we observed overall enrichment of regulatory and immune pathways in the top associated DNAm probes. Although previous studies have indicated that antipsychotic drugs may induce changes in DNA methylation (Burghardt et al., 2016; Houtepen et al., 2016; Kinoshita et al., 2017; Rukova et al., 2014), we did not find evidence for immediate large effects on DNA methylation based on CpG sites assayed in our experiments. Possibly, subtle changes in DNA methylation patterns play a regulatory role in the observed gene expression changes but a larger sample size is needed to decipher these changes(Jones, 2012). In addition, a wider range of clozapine concentrations *in vitro* may provide finer experimental resolution to observe subtle epigenetic changes and identify key regulatory drivers. DNA methylation is one type of epigenetic regulation and assaying other regulatory mechanisms, such as histone modification or RNA regulation, may provide further insights into the regulatory dynamics that drive widespread changes in gene expression (Allis and Jenuwein, 2016). Lastly, we observed DNA methylation changes to be time-dependent. While the clinical meaning of this remains speculative, our findings do suggest that the existing literature should be evaluated in the context of the duration of drug exposure. In addition, future studies could gain more insights into the function of clozapine when modeling drug exposure time as a variable in their analyses.

To further explore our gene expression findings, we investigated their association with genetic susceptibility of schizophrenia. We did not observe significant enrichment of schizophrenia heritability across clozapine-associated genes. Two previous studies did report an association between schizophrenia heritability and antipsychotic target genes(Gaspar and Breen, 2017; Skene et al., 2018). Their approach, however, differed from ours. While they used lists of antipsychotic target genes originating from pharmacological databases, we used a list of experimentally derived gene sets after *in vitro* exposure to clozapine in LCLs. The genes we found to be differentially expressed did not overlap with antipsychotic drug target genes used in these two previous studies, which may explain the discrepancy in findings. If we assume that the clozapine-induced gene expression differences are a proxy for adverse effects, the lack of evidence of the enrichment analysis suggests that the genetic architecture of disease susceptibility is likely independent from susceptibility to adverse effects. We explored this further by examining the enrichment of genetic signal linked to cholesterol and cardiovascular disease-related traits such as body mass index, coronary artery disease, type 2 diabetes, and triglyceride levels. We observed a nominally significant increased enrichment for total cholesterol and LDL genetic signal in upregulated genes in response to clozapine exposure. While these findings did not survive multiple testing correction and should be interpreted with caution, the observed cholesterol and LDL heritability enrichment increased as we narrowed down on the most strongly associated clozapine genes suggesting that this association warrants further investigation. Mapping heritability from large-scale population studies to *in vitro* experimental systems can then serve as a powerful approach to study biological pathways through integration of polygenic disease risk (Ori et al., 2019). Metabolic adverse effects are often observed in patients using clozapine (and antipsychotics in general) and improving our understanding of the mechanisms that underlie these adverse effects is an imperative area of research.

An important advantage of *in vitro* experimental models is the controlled laboratory environment, which not only decreases the signal-to-noise ratio in the collected data but also allows for precise manipulation of the model in follow up work. While our current results are consistent with previous findings in other cell types(Choi et al., 2009; Fernø et al., 2009, 2005; Hu et al., 2010; Lauressergues et al., 2012, 2011, 2010; Liu et al., 2009; Raeder et al., 2006; Vik-Mo et al., 2009; Yang et al., 2007, 2009), it remains unclear whether LCLs are an appropriate cell type to capture the molecular changes that are most relevant for studying adverse effects of antipsychotics in patients. To gain insight into the transferability of the *in vitro* clozapine-response in LCLs to functions of human tissues, we investigated how clozapine-associated genes are expressed across GTEx tissues. As gene expression patterns have been shown to have significant degree of sharing across human tissues(Buil et al., n.d.; GTEx Consortium et al., 2017) effects discovered in one tissue may thus be informative for other tissues. We observed that preferentially expressed genes in GTEx-LCL tissue overlap the most with clozapine genes identified in LCLs *in vitro*, with whole blood ranked as second highest tissue. Preferentially expressed genes however, represent only a subset of the clozapine genes. Using all associated genes, we demonstrated that upregulated clozapine genes have significantly different average expression in liver, muscle, lung, and testis tissue. This suggests multi-tissue downstream effects through clozapine treatment, which fits its clinical presentation of a diverse set of adverse effects that have been reported (Iqbal et al., 2003). Our finding of higher expression in the liver is in line with the observed cholesterol gene ontology signature of upregulated clozapine genes in our assay. The liver is a central player in the function of cholesterol in the body and clozapine-related hepatotoxicity has furthermore been reported in cases of treatment with the drug (Keane et al., 2009; Kellner et al., 1993). We observe no differences for any of the GTEx brain tissues.

On top of *within tissue* effects, we also performed a more explorative analysis to examine clozapine gene expression-based similarity *between tissues* and found specific tissue pairs that were significantly different in tissue similarities. Here, we highlight some interesting observations. We observe a decrease in similarity for tissues involved in immune (spleen), endocrine (testis, ovary, adrenal and thyroid gland) and metabolic (adipose) functioning. The spleen and testis in particular stand out with a significant change in dissimilarity with many tissues. The spleen is the largest secondary lymphoid organ in the body and hosts a wide range of immunological functions, including the storage of leukocytes (Lewis et al., 2019). While it remains speculative if splenic dysfunction could for example lead to agranulocytosis, our findings do solicit for more research on the mechanism between clozapine and splenic function. An intriguing, and perhaps also surprising, finding is the implication of the testis among analysis of genes downregulated by clozapine. The testis has two primary functions; to produce sperm and to produce hormones, in particular testosterone, which is a sex steroid synthesized from cholesterol (Eacker et al., 2008). Genes downregulated by clozapine are enriched for cell division processes and cell proliferation is an important function of testicular cells (Sohni et al., 2019). While little is currently known about how clozapine or other antipsychotics may affect testicular function in adult humans, our finding does again point to cholesterol-related biology. In addition to the testis, we also observe deviation in between-tissue similarity for several other endocrine tissues. As endocrine abnormalities are known causes of human obesity, the identified transcriptomic profile of clozapine response together with downstream tissue effects may point to new avenues to study clozapine-induced weight gain (Baptista, 1999). Together, these findings suggest that clozapine disproportionately affects the function of specific tissues. Our results furthermore demonstrate how experimental molecular signatures can be integrated with external genomic datasets, such as the GTEx project, to help translate in vitro findings to human biology.

We used LCLs as an *in vitro* model for studying molecular effects of clozapine exposure in an effort to improve our understanding of antipsychotic-induced adverse effects. Genome-wide gene expression profiling demonstrated a robust up-regulation of cholesterol metabolism and down-regulation of cell cycle pathways, with only limited changes in DNA methylation profiles. We did not find evidence that genes up-or down-regulated during clozapine exposure were enriched for genetic variation associated with schizophrenia. On the other hand, the observed enrichment signal with the genetic basis of total cholesterol and LDL levels and multi-tissue involved across immune, endocrine, and metabolic functions may provide important leads linked to antipsychotic drug induced metabolic adverse effects. The necessary challenge of large-scale, systematic prospective patient cohort studies of adverse effects of clozapine and other AP remains, while *in vitro* studies such as ours provide only glimpses of what may be relevant.

## Acknowledgements

The authors would like to thank Drs. Barbara Franke, Jeffrey Glennon, Jan Buitelaar, and Wouter Staal for their input in the early stages of the project and the GTEx Project and all used GWAS studies for making their data publicly available.

## References

1000 Genomes Project Consortium, Auton A, Brooks LD, Durbin RM, Garrison EP, Kang HM, Korbel JO, Marchini JL, McCarthy S, McVean GA, Abecasis GR. 2015. A global reference for human genetic variation. Nature 526:68–74.

Adkins DE, Aberg K, McClay JL, Bukszár J, Zhao Z, Jia P, Stroup TS, Perkins D, McEvoy JP, Lieberman JA, Sullivan PF, van den Oord EJCG. 2011. Genomewide pharmacogenomic study of metabolic side effects to antipsychotic drugs. Mol Psychiatry 16:321–332.

Allis CD, Jenuwein T. 2016. The molecular hallmarks of epigenetic control. Nat Rev Genet 17:487–500.

Andersohn F, Konzen C, Garbe E. 2007. Systematic review: agranulocytosis induced by nonchemotherapy drugs. Ann Intern Med 146:657–665.

Baptista T. 1999. Body weight gain induced by antipsychotic drugs: mechanisms and management. Acta Psychiatr Scand 100:3–16.

Baumann P, Hiemke C, Ulrich S, Eckermann G, Gaertner I, Gerlach M, Kuss H-J, Laux G, Müller-Oerlinghausen B, Rao ML, Riederer P, Zernig G, Arbeitsge-meinschaft fur neuropsychopharmakologie und pharmakopsychiatrie. 2004. The AGNP-TDM expert group consensus guidelines: therapeutic drug monitoring in psychiatry. Pharmacopsychiatry 37:243–265.

Buil A, Viñuela A, Brown AA, Davies MN, Padioleau I, Bielser D, Romano L, Glass D, Di Meglio P, Small KS, Spector TD, Dermitzakis ET. n.d. Quantifying the degree of sharing of genetic and non-genetic causes of gene expression variability across four tissues. doi: 10.1101/053355

Bulik-Sullivan B, Finucane HK, Anttila V, Gusev A, Day FR, Consortium R, Genomics Consortium P, of the Wellcome Trust Consortium GCFA, Perry JRB, Patterson N, Robinson E, Daly MJ, Price AL, Neale BM. 2015. An Atlas of Genetic Correlations across Human Diseases and Traits. Nat Genet 47:1237–1241.

Bulik-Sullivan BK, Loh P-R, Finucane HK, Ripke S, Yang J, Consortium SWG of TPG, Patterson N, Daly MJ, Price AL, Neale BM. 2015. LD Score regression distinguishes confounding from polygenicity in genome-wide association studies. Nat Genet 47:291–295.

Burghardt KJ, Goodrich JM, Dolinoy DC, Ellingrod VL. 2016. Gene-specific DNA methylation may mediate atypical antipsychotic-induced insulin resistance. Bipolar Disord 18:423–432.

Chen Y-A, Lemire M, Choufani S, Butcher DT, Grafodatskaya D, Zanke BW, Gallinger S, Hudson TJ, Weksberg R. 2013. Discovery of cross-reactive probes and polymorphic CpGs in the Illumina Infinium HumanMethylation450 microarray. Epigenetics 8:203–209.

Choi KH, Higgs BW, Weis S, Song J, Llenos IC, Dulay JR, Yolken RH, Webster MJ. 2009. Effects of typical and atypical antipsychotic drugs on gene expression profiles in the liver of schizophrenia subjects. BMC Psychiatry 9:57.

Chowdhury NI, Remington G, Kennedy JL. 2011. Genetics of antipsychotic-induced side effects and agranulocytosis. Curr Psychiatry Rep 13:156–165.

Cohen D. 2014. Prescribers fear as a major side-effect of clozapine. Acta Psychiatr Scand.

Consortium †the International Hapmap, †The International HapMap Consortium. 2003. The International HapMap Project. Nature. doi: 10.1038/nature02168

de Leeuw CA, Mooij JM, Heskes T, Posthuma D. 2015. MAGMA: Generalized Gene-Set Analysis of GWAS Data. PLoS Comput Biol 11. doi: 10.1371/journal.pcbi.1004219

de With SAJ, Pulit SL, Wang T, Staal WG, van Solinge WW, de Bakker PIW, Ophoff RA. 2015. Genome-wide association study of lymphoblast cell viability after clozapine exposure. Am J Med Genet B Neuropsychiatr Genet 168B:116–122.

Dietschy JM. 2009. Central nervous system: cholesterol turnover, brain development and neurodegeneration. Biological Chemistry. doi: 10.1515/bc.2009.035

Du P, Kibbe W a., Lin SM. 2008. lumi: a pipeline for processing Illumina microarray. Bioinformatics 24:1547–1548.

Du P, Zhang X, Huang C-C, Jafari N, Kibbe WA, Hou L, Lin SM. 2010. Comparison of Beta-value and M-value methods for quantifying methylation levels by microarray analysis. BMC Bioinformatics 11:587.

Eacker SM, Agrawal N, Qian K, Dichek HL, Gong E-Y, Lee K, Braun RE. 2008. Hormonal regulation of testicular steroid and cholesterol homeostasis. Mol Endocrinol 22:623–635.

Fernø J, Raeder MB, Vik-Mo AO, Skrede S, Glambek M, Tronstad K-J, Breilid H, Løvlie R, Berge RK, Stansberg C, Steen VM. 2005. Antipsychotic drugs activate SREBP-regulated expression of lipid biosynthetic genes in cultured human glioma cells: a novel mechanism of action? Pharmacogenomics J 5:298–304.

Ferno J, Skrede S, Vik-Mo AO, Jassim G, Le Hellard S, Steen VM. 2011. Lipogenic effects of psychotropic drugs: focus on the SREBP system. Front Biosci 16:49–60.

Fernø J, Vik-Mo AO, Jassim G, Håvik B, Berge K, Skrede S, Gudbrandsen OA, Waage J, Lunder N, Mørk S, Berge RK, Jørgensen HA, Steen VM. 2009. Acute clozapine exposure in vivo induces lipid accumulation and marked sequential changes in the expression of SREBP, PPAR, and LXR target genes in rat liver. Psychopharmacology 203:73–84.

Foley DL, Mackinnon A. 2014. A systematic review of antipsychotic drug effects on human gene expression related to risk factors for cardiovascular disease. Pharmacogenomics J 14:446–451.

Gaspar HA, Breen G. 2017. Drug enrichment and discovery from schizophrenia genome-wide association results: an analysis and visualisation approach. Sci Rep 7:12460.

Gassó P, Mas S, Rodríguez N, Boloc D, García-Cerro S, Bernardo M, Lafuente A, Parellada E. 2017. Microarray gene-expression study in fibroblast and lymphoblastoid cell lines from antipsychotic-naïve first-episode schizophrenia patients. J Psychiatr Res 95:91–101.

Gebhardt S, Theisen FM, Haberhausen M, Heinzel-Gutenbrunner M, Wehmeier PM, -C. Krieg J, Kühnau W, Schmidtke J, Remschmidt H, Hebebrand J. 2010. Body weight gain induced by atypical antipsychotics: an extension of the monocygotic twin and sib pair study. Journal of Clinical Pharmacy and Therapeutics. doi: 10.1111/j.1365-2710.2009.01084.x

GTEx Consortium, Laboratory, Data Analysis &Coordinating Center (LDACC)—Analysis Working Group, Statistical Methods groups—Analysis Working Group, Enhancing GTEx (eGTEx) groups, NIH Common Fund, NIH/NCI, NIH/NHGRI, NIH/NIMH, NIH/NIDA, Biospecimen Collection Source Site—NDRI, Biospecimen Collection Source Site—RPCI, Biospecimen Core Resource—VARI, Brain Bank Repository—University of Miami Brain Endowment Bank, Leidos Biomedical—Project Management, ELSI Study, Genome Browser Data Integration &Visualization—EBI, Genome Browser Data Integration &Visualization—UCSC Genomics Institute, University of California Santa Cruz, Lead analysts:, Laboratory, Data Analysis &Coordinating Center (LDACC):, NIH program management:, Biospecimen collection:, Pathology:, eQTL manuscript working group:, Battle A, Brown CD, Engelhardt BE, Montgomery SB. 2017. Genetic effects on gene expression across human tissues. Nature 550:204–213.

Harrison RNS, Murray RM, Lee SH, Paya Cano J, Dempster D, Curtis CJ, Dima D, Gaughran F, Breen G, de Jong S. 2016. Gene-expression analysis of clozapine treatment in whole blood of patients with psychosis. Psychiatr Genet 26:211–217.

Horvath S. 2011. Weighted Network Analysis: Applications in Genomics and Systems Biology. Springer Science & Business Media.

Houtepen LC, van Bergen AH, Vinkers CH, Boks MPM. 2016. DNA methylation signatures of mood stabilizers and antipsychotics in bipolar disorder. Epigenomics 8:197–208.

Huang DW, Lempicki R a., Sherman BT. 2009a. Systematic and integrative analysis of large gene lists using DAVID bioinformatics resources. Nat Protoc 4:44–57.

Huang DW, Sherman BT, Lempicki RA. 2009b. Bioinformatics enrichment tools: Paths toward the comprehensive functional analysis of large gene lists. Nucleic Acids Res 37:1–13.

Hu Y, Kutscher E, Davies GE. 2010. Berberine inhibits SREBP-1-related clozapine and risperidone induced adipogenesis in 3T3-L1 cells. Phytother Res 24:1831–1838.

Iqbal MM, Rahman A, Husain Z, Mahmud SZ, Ryan WG, Feldman JM. 2003. Clozapine: a clinical review of adverse effects and management. Ann Clin Psychiatry 15:33–48.

Jones PA. 2012. Functions of DNA methylation: islands, start sites, gene bodies and beyond. Nat Rev Genet 13:484–492.

Kane J, Honigfeld G, Singer J, Meltzer H. 1988. Clozapine for the treatment-resistant schizophrenic. A double-blind comparison with chlorpromazine. Arch Gen Psychiatry 45:789–796.

Keane S, Lane A, Larkin T, Clarke M. 2009. Management of clozapine-related hepatotoxicity. J Clin Psychopharmacol 29:606–607.

Kellner M, Wiedemann K, Krieg JC, Berg PA. 1993. Toxic hepatitis by clozapine treatment. Am J Psychiatry 150:985–986.

Kinoshita M, Numata S, Tajima A, Yamamori H, Yasuda Y, Fujimoto M, Watanabe S, Umehara H, Shimodera S, Nakazawa T, Kikuchi M, Nakaya A, Hashimoto H, Imoto I, Hashimoto R, Ohmori T. 2017. Effect of Clozapine on DNA Methylation in Peripheral Leukocytes from Patients with Treatment-Resistant Schizophrenia. Int J Mol Sci 18. doi: 10.3390/ijms18030632

Lahdelma L, Oja S, Korhonen M, Andersson LC. 2010. Clozapine is cytotoxic to primary cultures of human bone marrow mesenchymal stromal cells. J Clin Psychopharmacol 30:461–463.

Langfelder P, Horvath S. 2008. WGCNA: an R package for weighted correlation network analysis. BMC Bioinformatics 9:559.

Lauressergues E, Bert E, Duriez P, Hum D, Majd Z, Staels B, Cussac D. 2012. Does endoplasmic reticulum stress participate in APD-induced hepatic metabolic dysregulation? Neuropharmacology 62:784–796.

Lauressergues E, Martin F, Helleboid A, Bouchaert E, Cussac D, Bordet R, Hum D, Luc G, Majd Z, Staels B, Duriez P. 2011. Overweight induced by chronic risperidone exposure is correlated with overexpression of the SREBP-1c and FAS genes in mouse liver. Naunyn Schmiedebergs Arch Pharmacol 383:423–436.

Lauressergues E, Staels B, Valeille K, Majd Z, Hum DW, Duriez P, Cussac D. 2010. Antipsychotic drug action on SREBPs-related lipogenesis and cholesterogenesis in primary rat hepatocytes. Naunyn Schmiedebergs Arch Pharmacol 381:427–439.

Le Hellard S, Mühleisen TW, Djurovic S, Fernø J, Ouriaghi Z, Mattheisen M, Vasilescu C, Raeder MB, Hansen T, Strohmaier J, Georgi A, Brockschmidt FF, Melle I, Nenadic I, Sauer H, Rietschel M, Nöthen MM, Werge T, Andreassen OA, Cichon S, Steen VM. 2010. Polymorphisms in SREBF1 and SREBF2, two antipsychotic-activated transcription factors controlling cellular lipogenesis, are associated with schizophrenia in German and Scandinavian samples. Mol Psychiatry 15:463–472.

Lett TAP, Wallace TJM, Chowdhury NI, Tiwari AK, Kennedy JL, Müller DJ. 2012. Pharmacogenetics of antipsychotic-induced weight gain: review and clinical implications. Mol Psychiatry 17:242–266.

Leucht S, Cipriani A, Spineli L, Mavridis D, Orey D, Richter F, Samara M, Barbui C, Engel RR, Geddes JR, Kissling W, Stapf MP, Lässig B, Salanti G, Davis JM. 2013. Comparative efficacy and tolerability of 15 antipsychotic drugs in schizophrenia: a multiple-treatments meta-analysis. Lancet 382:951–962.

Lewis SM, Williams A, Eisenbarth SC. 2019. Structure and function of the immune system in the spleen. Sci Immunol 4. doi: 10.1126/sciimmunol.aau6085

Lin M, Pedrosa E, Hrabovsky A, Chen J, Puliafito BR, Gilbert SR, Zheng D, Lachman HM. 2016. Integrative transcriptome network analysis of iPSC-derived neurons from schizophrenia and schizoaffective disorder patients with 22q11.2 deletion. BMC Syst Biol 10:105.

Liu Y, Jandacek R, Rider T, Tso P, McNamara RK. 2009. Elevated delta-6 desaturase (FADS2) expression in the postmortem prefrontal cortex of schizophrenic patients: relationship with fatty acid composition. Schizophr Res 109:113–120.

Malhotra AK, Correll CU, Chowdhury NI, Müller DJ, Gregersen PK, Lee AT, Tiwari AK, Kane JM, Fleischhacker WW, Kahn RS, Ophoff RA, Meltzer HY, Lencz T, Kennedy JL. 2012. Association between common variants near the melanocortin 4 receptor gene and severe antipsychotic drug-induced weight gain. Arch Gen Psychiatry 69:904–912.

McLean CY, Bristor D, Hiller M, Clarke SL, Schaar BT, Lowe CB, Wenger AM, Bejerano G. 2010. GREAT improves functional interpretation of cis-regulatory regions. Nat Biotechnol 28:495–501.

Melas PA, Rogdaki M, Ösby U, Schalling M, Lavebratt C, Ekström TJ. 2012. Epigenetic aberrations in leukocytes of patients with schizophrenia: association of global DNA methylation with antipsychotic drug treatment and disease onset. FASEB J 26:2712–2718.

Melé M, Ferreira PG, Reverter F, DeLuca DS, Monlong J, Sammeth M, Young TR, Goldmann JM, Pervouchine DD, Sullivan TJ, Johnson R, Segrè AV, Djebali S, Niarchou A, GTEx Consortium, Wright FA, Lappalainen T, Calvo M, Getz G, Dermitzakis ET, Ardlie KG, Guigó R. 2015. Human genomics. The human transcriptome across tissues and individuals. Science 348:660–665.

Morag A, Kirchheiner J, Rehavi M, Gurwitz D. 2010. Human lymphoblastoid cell line panels: novel tools for assessing shared drug pathways. Pharmacogenomics 11:327–340.

Müller DJ, Chowdhury NI, Zai CC. 2013. The pharmacogenetics of antipsychotic-induced adverse events. Curr Opin Psychiatry 26:144–150.

Nikpay M, Goel A, Won H-H, Hall LM, Willenborg C, Kanoni S, Saleheen D, Kyriakou T, Nelson CP, Hopewell JC, Webb TR, Zeng L, Dehghan A, Alver M, Armasu SM, Auro K, Bjonnes A, Chasman DI, Chen S, Ford I, Franceschini N, Gieger C, Grace C, Gustafsson S, Huang J, Hwang S-J, Kim YK, Kleber ME, Lau KW, Lu X, Lu Y, Lyytikäinen L-P, Mihailov E, Morrison AC, Pervjakova N, Qu L, Rose LM, Salfati E, Saxena R, Scholz M, Smith AV, Tikkanen E, Uitterlinden A, Yang X, Zhang W, Zhao W, de Andrade M, de Vries PS, van Zuydam NR, Anand SS, Bertram L, Beutner F, Dedoussis G, Frossard P, Gauguier D, Goodall AH, Gottesman O, Haber M, Han B-G, Huang J, Jalilzadeh S, Kessler T, König IR, Lannfelt L, Lieb W, Lind L, Lindgren CM, Lokki M-L, Magnusson PK, Mallick NH, Mehra N, Meitinger T, Memon F-U-R, Morris AP, Nieminen MS, Pedersen NL, Peters A, Rallidis LS, Rasheed A, Samuel M, Shah SH, Sinisalo J, Stirrups KE, Trompet S, Wang L, Zaman KS, Ardissino D, Boerwinkle E, Borecki IB, Bottinger EP, Buring JE, Chambers JC, Collins R, Cupples LA, Danesh J, Demuth I, Elosua R, Epstein SE, Esko T, Feitosa MF, Franco OH, Franzosi MG, Granger CB, Gu D, Gudnason V, Hall AS, Hamsten A, Harris TB, Hazen SL, Hengstenberg C, Hofman A, Ingelsson E, Iribarren C, Jukema JW, Karhunen PJ, Kim B-J, Kooner JS, Kullo IJ, Lehtimäki T, Loos RJF, Melander O, Metspalu A, März W, Palmer CN, Perola M, Quertermous T, Rader DJ, Ridker PM, Ripatti S, Roberts R, Salomaa V, Sanghera DK, Schwartz SM, Seedorf U, Stewart AF, Stott DJ, Thiery J, Zalloua PA, O’Donnell CJ, Reilly MP, Assimes TL, Thompson JR, Erdmann J, Clarke R, Watkins H, Kathiresan S, McPherson R, Deloukas P, Schunkert H, Samani NJ, Farrall M. 2015. A comprehensive 1,000 Genomes-based genome-wide association meta-analysis of coronary artery disease. Nat Genet 47:1121–1130.

Okazaki S, Boku S, Otsuka I, Mouri K, Aoyama S, Shiroiwa K, Sora I, Fujita A, Shirai Y, Shirakawa O, Kokai M, Hishimoto A. 2016. The cell cycle-related genes as biomarkers for schizophrenia. Prog Neuropsychopharmacol Biol Psychiatry 70:85–91.

Ori APS, Bot MHM, Molenhuis RT, Olde Loohuis LM, Ophoff RA. 2019. A Longitudinal Model of Human Neuronal Differentiation for Functional Investigation of Schizophrenia Polygenic Risk. Biol Psychiatry 85:544–553.

Ota VK, Noto C, Gadelha A, Santoro ML, Spindola LM, Gouvea ES, Stilhano RS, Ortiz BB, Silva PN, Sato JR, Han SW, Cordeiro Q, Bressan RA, Belangero SI. 2014. Changes in gene expression and methylation in the blood of patients with first-episode psychosis. Schizophr Res 159:358–364.

Pereira A, Dean B. 2006. Clozapine bioactivation induces dose-dependent, drug-specific toxicity of human bone marrow stromal cells: A potential in vitro system for the study of agranulocytosis. Biochemical Pharmacology. doi: 10.1016/j.bcp.2006.06.006

Pidsley R, Y Wong CC, Volta M, Lunnon K, Mill J, Schalkwyk LC. 2013. A data-driven approach to preprocessing Illumina 450K methylation array data. BMC Genomics 14:293.

Price ME, Cotton AM, Lam LL, Farré P, Emberly E, Brown CJ, Robinson WP, Kobor MS. 2013. Additional annotation enhances potential for biologically-relevant analysis of the Illumina Infinium HumanMethylation450 BeadChip array. Epigenetics Chromatin 6:4.

Raeder MB, Fernø J, Vik-Mo AO, Steen VM. 2006. SREBP activation by antipsychotic- and antidepressant-drugs in cultured human liver cells: relevance for metabolic side-effects? Mol Cell Biochem 289:167–173.

Roerig JL, Steffen KJ, Mitchell JE. 2011. Atypical antipsychotic-induced weight gain: insights into mechanisms of action. CNS Drugs 25:1035–1059.

Roth BL, Lopez E, Patel S, Kroeze WK. 2000. The Multiplicity of Serotonin Receptors: Uselessly Diverse Molecules or an Embarrassment of Riches? The Neuroscientist. doi: 10.1177/107385840000600408

Rukova B, Staneva R, Hadjidekova S, Stamenov G, Milanova V, Toncheva D. 2014. Whole genome methylation analyses of schizophrenia patients before and after treatment. Biotechnol Biotechnol Equip 28:518–524.

Schizophrenia Working Group of the Psychiatric Genomics Consortium. 2014. Biological insights from 108 schizophrenia-associated genetic loci. Nature. doi: 10.1038/nature13595

Scott RA, Scott LJ, Mägi R, Marullo L, Gaulton KJ, Kaakinen M, Pervjakova N, Pers TH, Johnson AD, Eicher JD, Jackson AU, Ferreira T, Lee Y, Ma C, Steinthorsdottir V, Thorleifsson G, Qi L, Van Zuydam NR, Mahajan A, Chen H, Almgren P, Voight BF, Grallert H, Müller-Nurasyid M, Ried JS, Rayner NW, Robertson N, Karssen LC, van Leeuwen EM, Willems SM, Fuchsberger C, Kwan P, Teslovich TM, Chanda P, Li M, Lu Y, Dina C, Thuillier D, Yengo L, Jiang L, Sparso T, Kestler HA, Chheda H, Eisele L, Gustafsson S, Frånberg M, Strawbridge RJ, Benediktsson R, Hreidarsson AB, Kong A, Sigurðsson G, Kerrison ND, Luan J ’an, Liang L, Meitinger T, Roden M, Thorand B, Esko T, Mihailov E, Fox C, Liu C-T, Rybin D, Isomaa B, Lyssenko V, Tuomi T, Couper DJ, Pankow JS, Grarup N, Have CT, Jørgensen ME, Jørgensen T, Linneberg A, Cornelis MC, van Dam RM, Hunter DJ, Kraft P, Sun Q, Edkins S, Owen KR, Perry JRB, Wood AR, Zeggini E, Tajes-Fernandes J, Abecasis GR, Bonnycastle LL, Chines PS, Stringham HM, Koistinen HA, Kinnunen L, Sennblad B, Mühleisen TW, Nöthen MM, Pechlivanis S, Baldassarre D, Gertow K, Humphries SE, Tremoli E, Klopp N, Meyer J, Steinbach G, Wennauer R, Eriksson JG, Männistö S, Peltonen L, Tikkanen E, Charpentier G, Eury E, Lobbens S, Gigante B, Leander K, McLeod O, Bottinger EP, Gottesman O, Ruderfer D, Blüher M, Kovacs P, Tonjes A, Maruthur NM, Scapoli C, Erbel R, Jöckel K-H, Moebus S, de Faire U, Hamsten A, Stumvoll M, Deloukas P, Donnelly PJ, Frayling TM, Hattersley AT, Ripatti S, Salomaa V, Pedersen NL, Boehm BO, Bergman RN, Collins FS, Mohlke KL, Tuomilehto J, Hansen T, Pedersen O, Barroso I, Lannfelt L, Ingelsson E, Lind L, Lindgren CM, Cauchi S, Froguel P, Loos RJF, Balkau B, Boeing H, Franks PW, Barricarte Gurrea A, Palli D, van der Schouw YT, Altshuler D, Groop LC, Langenberg C, Wareham NJ, Sijbrands E, van Duijn CM, Florez JC, Meigs JB, Boerwinkle E, Gieger C, Strauch K, Metspalu A, Morris AD, Palmer CNA, Hu FB, Thorsteinsdottir U, Stefansson K, Dupuis J, Morris AP, Boehnke M, McCarthy MI, Prokopenko I, DIAbetes Genetics Replication and Meta-analysis (DIAGRAM) Consortium. 2017. An Expanded Genome-Wide Association Study of Type 2 Diabetes in Europeans. Diabetes 66:2888–2902.

Skene NG, Bryois J, Bakken TE, Breen G, Crowley JJ, Gaspar HA, Giusti-Rodriguez P, Hodge RD, Miller JA, Muñoz-Manchado AB, O’Donovan MC, Owen MJ, Pardiñas AF, Ryge J, Walters JTR, Linnarsson S, Lein ES, Major Depressive Disorder Working Group of the Psychiatric Genomics Consortium, Sullivan PF, Hjerling-Leffler J. 2018. Genetic identification of brain cell types underlying schizophrenia. Nat Genet 50:825–833.

Smyth GK. n.d. limma: Linear Models for Microarray Data. Bioinformatics and Computational Biology Solutions Using R and Bioconductor. doi: 10.1007/0-387-29362-0_23

Sohni A, Tan K, Song H-W, Burow D, de Rooij DG, Laurent L, Hsieh T-C, Rabah R, Hammoud SS, Vicini E, Wilkinson MF. 2019. The Neonatal and Adult Human Testis Defined at the Single-Cell Level. Cell Rep 26:1501–1517.e4.

Stapel B, Kotsiari A, Scherr M, Hilfiker-Kleiner D, Bleich S, Frieling H, Kahl KG. 2017. Olanzapine and aripiprazole differentially affect glucose uptake and energy metabolism in human mononuclear blood cells. J Psychiatr Res 88:18–27.

Stroup TS, Gerhard T, Crystal S, Huang C, Olfson M. 2016. Comparative Effectiveness of Clozapine and Standard Antipsychotic Treatment in Adults With Schizophrenia. Am J Psychiatry 173:166–173.

Surakka I, Horikoshi M, Mägi R, Sarin A-P, Mahajan A, Lagou V, Marullo L, Ferreira T, Miraglio B, Timonen S, Kettunen J, Pirinen M, Karjalainen J, Thorleifsson G, Hägg S, Hottenga J-J, Isaacs A, Ladenvall C, Beekman M, Esko T, Ried JS, Nelson CP, Willenborg C, Gustafsson S, Westra H-J, Blades M, de Craen AJM, de Geus EJ, Deelen J, Grallert H, Hamsten A, Havulinna AS, Hengstenberg C, Houwing-Duistermaat JJ, Hyppönen E, Karssen LC, Lehtimäki T, Lyssenko V, Magnusson PKE, Mihailov E, Müller-Nurasyid M, Mpindi J-P, Pedersen NL, Penninx BWJH, Perola M, Pers TH, Peters A, Rung J, Smit JH, Steinthorsdottir V, Tobin MD, Tsernikova N, van Leeuwen EM, Viikari JS, Willems SM, Willemsen G, Schunkert H, Erdmann J, Samani NJ, Kaprio J, Lind L, Gieger C, Metspalu A, Slagboom PE, Groop L, van Duijn CM, Eriksson JG, Jula A, Salomaa V, Boomsma DI, Power C, Raitakari OT, Ingelsson E, Järvelin M-R, Thorsteinsdottir U, Franke L, Ikonen E, Kallioniemi O, Pietiäinen V, Lindgren CM, Stefansson K, Palotie A, McCarthy MI, Morris AP, Prokopenko I, Ripatti S, ENGAGE Consortium. 2015. The impact of low-frequency and rare variants on lipid levels. Nat Genet 47:589–597.

Swathy B, Banerjee M. 2017. Haloperidol induces pharmacoepigenetic response by modulating miRNA expression, global DNA methylation and expression profiles of methylation maintenance genes and genes involved in neurotransmission in neuronal cells. PLoS One 12:e0184209.

Swathy B, Saradalekshmi KR, Nair IV, Nair C, Banerjee M. 2017. Pharmacoepigenomic responses of antipsychotic drugs on pharmacogenes are likely to be modulated by miRNAs. Epigenomics 9:811–821.

Taylor DM. 2017. Clozapine for Treatment-Resistant Schizophrenia: Still the Gold Standard? CNS Drugs 31:177–180.

Venables WN, Ripley BD. 2002. Modern Applied Statistics with S. Statistics and Computing. doi: 10.1007/978-0-387-21706-2

Vik-Mo AO, Birkenaes AB, Fernø J, Jonsdottir H, Andreassen OA, Steen VM. 2008. Increased expression of lipid biosynthesis genes in peripheral blood cells of olanzapine-treated patients. Int J Neuropsychopharmacol 11:679–684.

Vik-Mo AO, Fernø J, Skrede S, Steen VM. 2009. Psychotropic drugs up-regulate the expression of cholesterol transport proteins including ApoE in cultured human CNS- and liver cells. BMC Pharmacol 9:10.

Wagner AH, Coffman AC, Ainscough BJ, Spies NC, Skidmore ZL, Campbell KM, Krysiak K, Pan D, McMichael JF, Eldred JM, Walker JR, Wilson RK, Mardis ER, Griffith M, Griffith OL. 2016. DGIdb 2.0: mining clinically relevant drug–gene interactions. Nucleic Acids Research. doi: 10.1093/nar/gkv1165

Wang L, Lockstone HE, Guest PC, Levin Y, Palotás A, Pietsch S, Schwarz E, Rahmoune H, Harris LW, Ma D, Bahn S. 2010. Expression profiling of fibroblasts identifies cell cycle abnormalities in schizophrenia. J Proteome Res 9:521–527.

Weiden PJ, Mackell JA, McDonnell DD. 2004. Obesity as a risk factor for antipsychotic noncompliance. Schizophrenia Research. doi: 10.1016/s0920-9964(02)00498-x

Welsh M, Mangravite L, Medina MW, Tantisira K, Zhang W, Huang RS, McLeod H, Dolan ME. 2009. Pharmacogenomic discovery using cell-based models. Pharmacol Rev 61:413–429.

Wen Y, Gamazon ER, Bleibel WK, Wing C, Mi S, McIlwee BE, Delaney SM, Duan S, Im HK, Dolan ME. 2012. An eQTL-based method identifies CTTN and ZMAT3 as pemetrexed susceptibility markers. Hum Mol Genet 21:1470–1480.

Williams DP, Pirmohamed M, Naisbitt DJ, Maggs JL, Park BK. 1997. Neutrophil cytotoxicity of the chemically reactive metabolite(s) of clozapine: possible role in agranulocytosis. J Pharmacol Exp Ther 283:1375–1382.

Yang L, Chen J, Liu D, Yu S, Cong E, Li Y, Wu H, Yue Y, Zuo S, Wang Y, Liang S, Shi Y, Shi S, Xu Y. 2015. Association between SREBF2 gene polymorphisms and metabolic syndrome in clozapine-treated patients with schizophrenia. Prog Neuropsychopharmacol Biol Psychiatry 56:136–141.

Yang L, Chen J, Li Y, Wang Y, Liang S, Shi Y, Shi S, Xu Y. 2016. Association between SCAP and SREBF1 gene polymorphisms and metabolic syndrome in schizophrenia patients treated with atypical antipsychotics. World J Biol Psychiatry 17:467–474.

Yang L-H, Chen T-M, Yu S-T, Chen Y-H. 2007. Olanzapine induces SREBP-1-related adipogenesis in 3T3-L1 cells. Pharmacol Res 56:202–208.

Yang Z, Li M, Hu X, Xiang B, Deng W, Wang Q, Wang Y, Zhao L, Ma X, Sham PC, Northoff G, Li T. 2017. Rare damaging variants in DNA repair and cell cycle pathways are associated with hippocampal and cognitive dysfunction: a combined genetic imaging study in first-episode treatment-naive patients with schizophrenia. Transl Psychiatry 7:e1028.

Yang Z, Yin J-Y, Gong Z-C, Huang Q, Chen H, Zhang W, Zhou H-H, Liu Z-Q. 2009. Evidence for an effect of clozapine on the regulation of fat-cell derived factors. Clin Chim Acta 408:98–104.

Yan H, Chen J-D, Zheng X-Y. 2013. Potential mechanisms of atypical antipsychotic-induced hypertriglyceridemia. Psychopharmacology 229:1–7.

Yengo L, Sidorenko J, Kemper KE, Zheng Z, Wood AR, Weedon MN, Frayling TM, Hirschhorn J, Yang J, Visscher PM, the GIANT Consortium. n.d. Meta-analysis of genome-wide association studies for height and body mass index in ∼700,000 individuals of European ancestry. doi: 10.1101/274654

Zhang B, Horvath S. 2005. A general framework for weighted gene co-expression network analysis. Stat Appl Genet Mol Biol 4:Article17.

