## Supplementary Materials for "Integrative genomic strategies applied to a lymphoblast cell line model reveal specific transcriptomic signatures associated with clozapine response"

**Table of contents:**

Supplementary Material and Methods 2
Clozapine exposure and in vitro experimental design 2
Sample processing and gene expression data 2
Sample processing and DNA methylation data 3
Gene ontology analysis 3
Functional enrichment analysis of DNA methylation data 3
Supplementary Figures 4
Supplementary Tables 6
Supplementary Results 7
Clozapine-associated genes and their preferential expression in GTEx tissues 7
References 8

### Supplementary Material and Methods

*Clozapine exposure and in vitro experimental design*

We performed drug exposure experiments in 6-well plates (Genesee, San Diego, CA, USA) and assessed cell viability using the TC10 automated cell counter (Bio-Rad, Hercules, CA, USA), according to manufacturer's instructions. Clozapine was purchased from Sigma Aldrich (St. Louis, MO, USA). Previous work has suggested that clinical concentrations of antipsychotics may not induce significant gene expression changes *in vitro* [1]. There is evidence that *in vivo* concentrations of antipsychotic drugs are higher in brain tissue than in peripheral blood [2, 3]. We therefore chose to expose cell lines to different clozapine concentrations, with clinical concentration set at 2μM [4]. Clozapine was dissolved in culture medium with dimethyl sulfoxide (DMSO), with a maximum concentration of 0.1%. Cell lines were exposed for 24 hours to clinical concentration, 10x, 50x and 100x clinical concentration (20μM-100 μM-200 μM clozapine) and vehicle (DMSO); each concentration was measured in 4 cell lines, after which RNA was obtained for gene expression analysis.

To study DNA methylation changes in response to clozapine, we performed an experiment similar to the gene expression study; LCLs were exposed to different concentrations of clozapine (vehicle (DMSO), 1x, 20x, 40x and 60 times clinical concentration) and exposure times were 24h and 96h (Supplemental Figure 1B).

*Sample processing and gene expression data*

We performed RNA extraction with Qiagen RNeasy mini kit (Qiagen, Valencia, CA, USA), according to manufacturer's instructions. RNA quantity and quality were measured with T2100 BioAnalyzer (Agilent, Santa Clara, CA, USA) and verified with a NanoDrop Spectrophotometer (NanoDrop products, Wilmington, DE, USA). Gene expression profiling was carried out using Illumina® HumanHT-12 v4 Expression BeadChip technology (Illumina, San Diego, CA, USA).

*Sample processing and DNA methylation data*

After desired exposure time, cells were lysed and DNA was extracted with DNeasy® Blood and Tissue kit (Qiagen, Valencia, CA, USA), according to the instructions of the manufacturer. DNA quality and quantity were assessed with the picogreen® assay (VWR, West Chester, PA, USA) and Nanodrop (ThermoScientific, Wilmington, DE, USA). DNA methylation assays were performed with Illumina® Infinium HumanMethylation450 Beadchip arrays (Illumina, San Diego, CA, USA), assaying approximately 450,000 CpG sites.

*Gene ontology analysis*

We performed functional gene ontology analysis using DAVID (Database for Annotation, Visualization and Integrated Discovery, version 6.8, interrogated February 2018) [5, 6], with default settings.

*Functional enrichment analysis of DNA methylation data*

Genomic Regions Enrichment of Annotations Tools (GREAT, v3.0) was used to predict the biological function of the top methylation probes associated to clozapine exposure. GREAT links both proximal and distal genomic CpG sites with their putative target genes and implements both a gene-based test and a region-based test using the hypergeometric and binomial test, respectively, to assess enrichment of genomic regions in biological annotations [7]. CpG sites were uploaded to the GREAT web portal (http://great.stanford.edu/public/html/) and analyses were run using the hg19 reference annotation and the whole genome as background. Genomic regions were assigned to genes if they are between 5 Kb upstream and 1 Kb downstream of the TSS, plus up to 1 Mb distal. Pathway annotations from GO Biological Processes, GO Cellular Component, GO Molecular Function, MSigDB, and PANTHER were used to infer biological meaning for CpG sites associated with clozapine.

### Supplementary Figures

Supplementary Figure 1A. Flow chart of gene expression analyses plan.

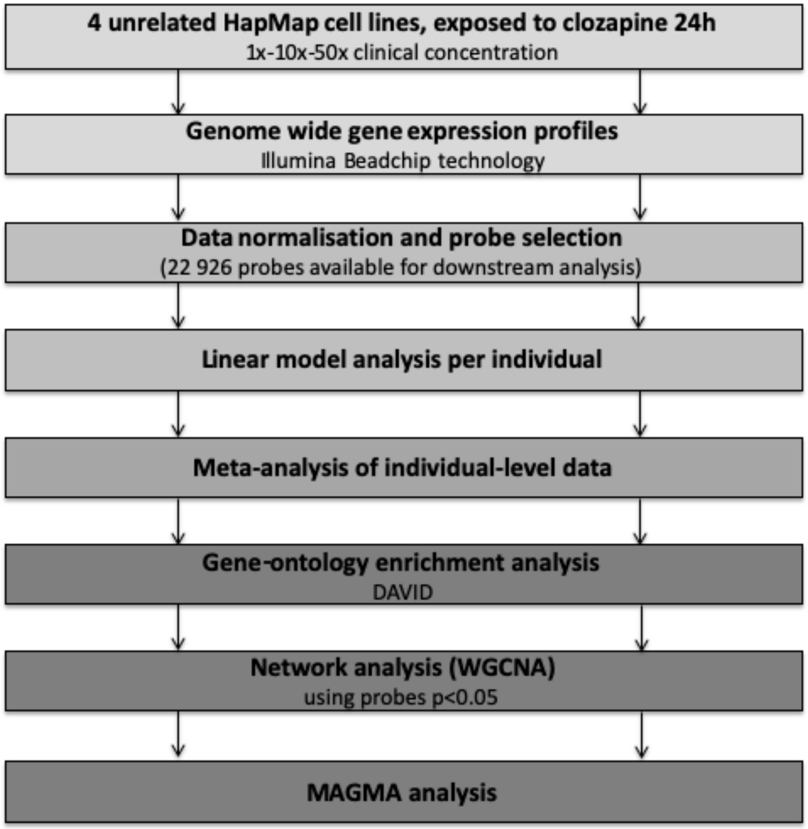

Supplementary Figure 1B. Flow chart of DNA methylation analyses plan.

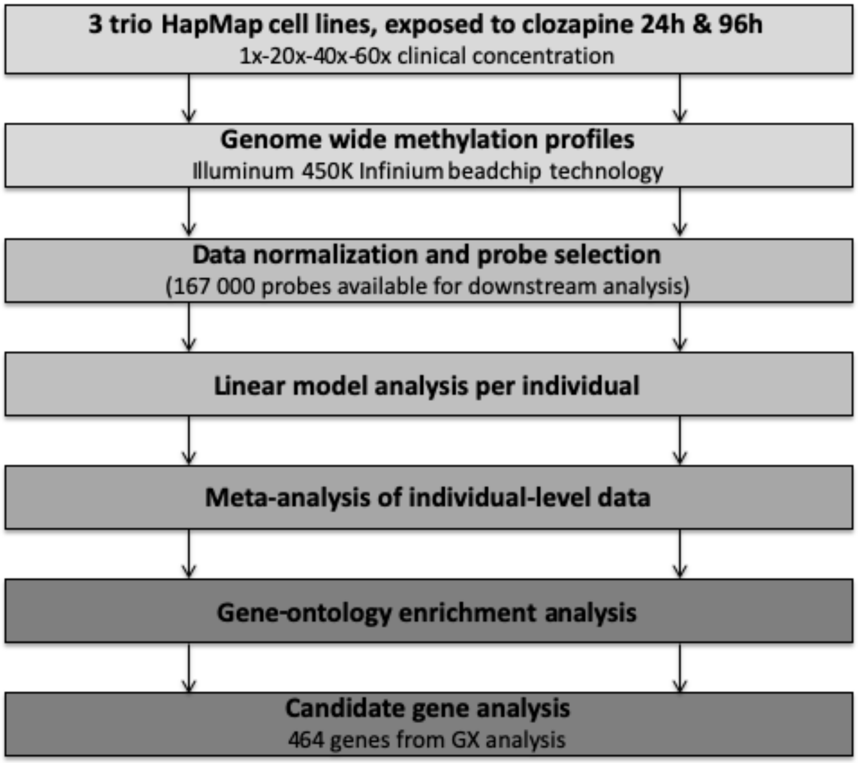

### Supplementary Tables

**Supplementary Table 1. List of differentially expressed genes**

Available as excel sheet.

**Supplementary Table 2. Top 10 DNA methylation probes after 24h and 96h of clozapine exposure.**

| 24h clozapine exposure | | | | | 96h clozapine exposure | | | | |
|---|---|---|---|---|---|---|---|---|---|
| Probe ID | Chr# | Coordinate | Annotation | P-value* | Probe ID | Chr# | Coordinate | Annotation | P-value* |
| cg09495017 | 16 | 57,124,763 | *CNOT1* | $5.40^{*}10^{-7}$ | cg21293934 | 18 | 14,738,230 | *ANKRD30B* | $8.87^{*}10^{-7}$ |
| cg16258062 | 2 | 234,048,887 | - | $5.60^{*}10^{-7}$ | cg19182557 | 2 | 130,061,825 | - | $9.21^{*}10^{-7}$ |
| cg16258062 | 1 | 47,654,324 | *FOXE3* | $1.15^{*}10^{-6}$ | cg15463280 | 11 | 95,955,596 | - | $2.36^{*}10^{-6}$ |
| cg15066636 | 6 | 33,187,127 | *HLA-DPB2* | $1.42^{*}10^{-6}$ | cg12564567 | 11 | 115,876,398 | - | $5.87^{*}10^{-6}$ |
| cg17488052 | 1 | 77,993,925 | *USP33* | $1.48^{*}10^{-6}$ | cg26647200 | 16 | 2,422,776 | *CCNF* | $6.87^{*}10^{-6}$ |
| cg25181236 | 4 | 56,082,032 | *CLOCK* | $1.48^{*}10^{-6}$ | cg09333631 | 3 | 44,777,608 | *KIF15, KIAA1143* | $7.99^{*}10^{-6}$ |
| cg01531409 | 14 | 59,781,022 | *PPM1A* | $2.07^{*}10^{-6}$ | cg16924010 | 3 | 195,500,852 | - | $8.51^{*}10^{-6}$ |
| cg09840472 | 7 | 22,730,922 | - | $2.27^{*}10^{-6}$ | cg01842314 | 10 | 106,102,325 | *CCDC147* | $1.07^{*}10^{-5}$ |
| cg24207009 | 17 | 73,549,157 | *TNRC6C* | $2.37^{*}10^{-6}$ | cg23898204 | 2 | 724,927 | - | $1.11^{*}10^{-5}$ |
| cg27170003 | 17 | 3,713,677 | *CAMKK1* | $2.39^{*}10^{-6}$ | cg15000279 | 19 | 33,976,849 | - | $1.28^{*}10^{-5}$ |

Supplementary Table 3. Candidate gene analysis: significant methylation probes after 24h and 96h of clozapine exposure.

| 24h clozapine exposure | | | | |
|---|---|---|---|---|
| Probe ID | Annotation | P-value | Gene expression Probe ID | Gene expression P-value |
| cg05455234 | *PCNT* *(Pericentrin)* | $1.98^{*}10^{-5}$ | ILMN_1810922 | $5.68^{*}10^{-8}$ |
| cg22971501 | *LDLR* *(Low Density Lipoprotein Receptor)* | $4.75^{*}10^{-5}$ | ILMN_2053415 | $3.97^{*}10^{-14}$ |
| cg01233620 | *CLEC16A* *(C-Type Lectin Domain Containing 16A)* | $6.30^{*}10^{-5}$ | ILMN_1781752 | $1.04^{*}10^{-7}$ |
| 96h clozapine exposure | | | | |
| cg26647200 | *CCNF* *(Cyclin F)* | $6.87^{*}10^{-6}$ | ILMN_1773119 | $4.41^{*}10^{-11}$ |

**Supplementary Table 8. Gene-set analyses results of the schizophrenia GWAS at varying significance levels.**

| Set | Bonferroni | FDR < 1% (q<0.01) | FDR<5% (q<0.05) |
|---|---|---|---|
| Differentially expressed genes | p = 0.92 (311 genes) | p = 0.74 (919 genes) | p = 0.22 (1543 genes) |
| Upregulated genes | p = 0.75 (138 genes) | p = 0.71 (457 genes) | p = 0.99 (772 genes) |
| Downregulated genes | p = 0.64 (173 genes) | p = 0.39 (462 genes) | p = 0.09 (771 genes) |

### Supplementary Results

*Clozapine-associated genes and their preferential expression in GTEx tissues*

To investigate tissue-specificity of genes identified to be differentially expressed after clozapine exposure, we used an available list of genes that are preferentially expressed in an individual tissue as identified by GTEx [8]. Preferential tissue expression is defined as all instances where the mean expression of the gene in the tested tissue was significantly higher (FDR<0.01 and a log2 fold change >= 4) than in the samples from the rest of the tissues. Visualized below is the fraction of genes with tissue preferential expression that are detected in our assay (x-axis) versus detected in our assay *and* associated to clozapine exposure (FDR < 5%) for each tissue. We observe that almost half of the differentially expressed genes have preferential expression in LCL tissue in GTEx. The second highest tissue is whole blood.

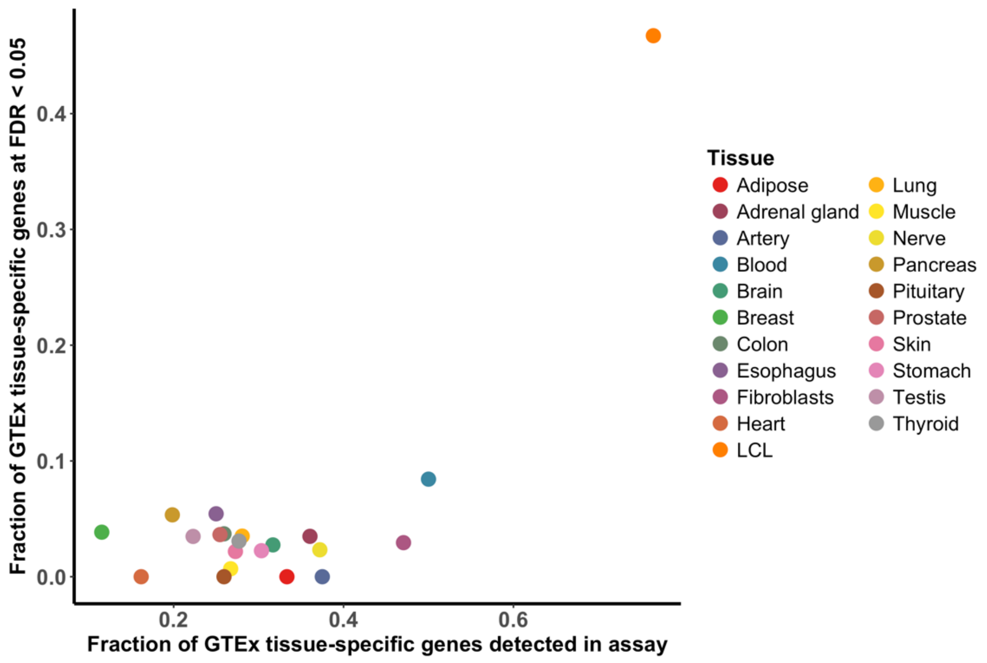


**Supplementary Figure 2. Differentially expressed genes are preferentially expressed in LCL tissue in GTEx.**

### References


1. Fernø J, Skrede S, Vik-Mo AO, Håvik B, Steen VM. Drug-induced activation of SREBP-controlled lipogenic gene expression in CNS-related cell lines: marked differences between various antipsychotic drugs. BMC Neurosci. 2006;7:69.
2. Weigmann H, Härtter S, Fischer V, Dahmen N, Hiemke C. Distribution of clozapine and desmethylclozapine between blood and brain in rats. Eur Neuropsychopharmacol. 1999;9:253–256.
3. Kornhuber J, Schultz A, Wiltfang J, Meineke I, Gleiter CH, Zöchling R, et al. Persistence of haloperidol in human brain tissue. Am J Psychiatry. 1999;156:885–890.
4. Baumann P, Hiemke C, Ulrich S, Eckermann G, Gaertner I, Gerlach M, et al. The AGNP-TDM expert group consensus guidelines: therapeutic drug monitoring in psychiatry. Pharmacopsychiatry. 2004;37:243–265.
5. Huang DW, Sherman BT, Lempicki RA. Bioinformatics enrichment tools: Paths toward the comprehensive functional analysis of large gene lists. Nucleic Acids Res. 2009;37:1–13.
6. Huang DW, Lempicki R a., Sherman BT. Systematic and integrative analysis of large gene lists using DAVID bioinformatics resources. Nat Protoc. 2009;4:44–57.
7. McLean CY, Bristor D, Hiller M, Clarke SL, Schaar BT, Lowe CB, et al. GREAT improves functional interpretation of cis-regulatory regions. Nat Biotechnol. 2010;28:495–501.
8. Melé M, Ferreira PG, Reverter F, DeLuca DS, Monlong J, Sammeth M, et al. Human genomics. The human transcriptome across tissues and individuals. Science. 2015;348:660–665.
